# Brain dysfunction during warming is caused by oxygen limitation in larval zebrafish

**DOI:** 10.1101/2020.12.28.424529

**Authors:** Anna H. Andreassen, Petter Hall, Pouya Khatibzadeh, Fredrik Jutfelt, Florence Kermen

**Author notes:** Corresponding author: Florence Kermen.

## Abstract

Understanding the physiological mechanisms that limit animal thermal tolerance is crucial in predicting how animals will respond to increasingly severe heatwaves. Despite its importance for understanding climate change impacts, these mechanisms underlying the upper thermal tolerance limits of animals are largely unknown. It has been hypothesised that the upper thermal tolerance in fish is limited by the thermal tolerance of the brain and that it is ultimately caused by a global brain depolarization. In this study, we developed methods for measuring the upper thermal limit (CT_max_) in larval zebrafish (*Danio rerio*) with simultaneous recordings of brain activity using GCaMP6s calcium imaging in both free-swimming and agar-embedded fish. We discovered that during warming, CT_max_ precedes, and is therefore not caused by, a global brain depolarization. Instead, the CT_max_ coincides with a decline in spontaneous neural activity and a loss of neural response to visual stimuli. By manipulating water oxygen levels, we found that oxygen availability during heating affects both locomotor-related neural activity, the neural response to visual stimuli, and CT_max_. Our results suggest that the mechanism limiting the upper thermal tolerance in zebrafish larvae is reduced oxygen availability causing impaired brain function.

## INTRODUCTION

Heatwaves are becoming more frequent and severe as climate change progresses (1, 2). Ectotherms have a similar body temperature to the surrounding environment and might be particularly vulnerable to extreme heat as their biological rates are directly affected by the ambient temperature (3–5). The range of temperatures an ectotherm can tolerate, is bounded at the upper end by a critical upper thermal limit (CT_max_), the temperature at which their locomotor function is lost and their movement becomes disorganized (6–8). The thermal tolerance limits of aquatic ectotherms correlates with the water temperature in geographical distributions (9), suggesting that limits in thermal tolerance partly determines distribution ranges. Furthermore, rapid water warming during heatwaves can cause mass mortality in fishes (10). Knowledge of the physiological mechanisms that control thermal tolerance is therefore an important part of understanding how ectotherms may respond to increasingly warmer temperatures and in predicting ecosystem impacts. Several hypotheses on the limitations of thermal tolerance have been proposed but the causes of thermal limits remains debated (11–13).

The hypothesis of oxygen limitation (i.e. the oxygen- and capacity-limited thermal tolerance (OCLTT) hypothesis) suggests that tissue oxygen shortage limits thermal tolerance under acute heat stress (14–16). According to this hypothesis, the cardiorespiratory system in ectotherms is unable to compensate for the temperature-induced increase in oxygen requirement, leading to tissue hypoxia and thus setting the limits for thermal tolerance. Insufficient cardiorespiratory function at high temperatures, or near upper thermal limits, has been reported in invertebrates and fishes (17–20), while this function is maintained in a number of other ectotherms (21, 22). In addition, experimental manipulation of oxygen availability during heat stress often fails to affect thermal tolerance (23). These contrasting results from a considerable amount of literature suggest that oxygen limitation can only partly explain thermal tolerance, and that these oxygen effects on thermal tolerance is species- and context-dependent (12, 16, 22, 24, 25). Lastly, the OCLTT hypothesis does not specify which physiological mechanisms fail first as a result of tissue hypoxia (24).

Another major hypothesis on limits of upper thermal tolerance is based on temperature-induced neural dysfunction. Neural control of locomotion is a heat-sensitive physiological mechanism that is disrupted at high temperatures and controlled locomotion is impaired at CT_max_. Temperature-induced neural dysfunction could consequently underlie upper thermal tolerance limits, either via direct thermal effects on neurons (11, 12) or via indirect thermal effects from oxygen limitations (18). Examples of severe heat-induced neural dysfunctions include spreading depolarizations in the brain of fruit flies (26), loss of rhythmic neural activity in the digestive system of the Jonah crab (27), and thermogenic seizures in vertebrates (28–30). However, few studies have directly investigated how neural dysfunction relates to measurements of thermal tolerance. In fruit flies, spreading depolarizations measured in restrained flies occur at similar temperatures as a heat-induced coma in freely moving conspecifics (26). Similarly, goldfish lose equilibrium at temperatures that alters the activity of cerebellar neurons in anesthetized conspecifics (11), and a study on Atlantic cod found that cooling the brain marginally increased CT_max_, suggesting a causal link between brain function and thermal tolerance (12).

The relative importance of a cardiorespiratory limitation versus direct thermal effects on neural function has been subject to considerable discussion in the literature on thermal limits (12, 24, 31–33). The investigation of thermal limits in neural function has been hindered by technical challenges. For example, it has not been possible to record brain activity and measure CT_max_ simultaneously during acute warming in freely moving animals.

In this study, we solved that challenge through a series of five heat ramping experiments on transparent larval zebrafish (*Danio rerio*) expressing a genetically encoded calcium indicator in neurons throughout the brain (34). This allowed simultaneous recording of neural activity, CT_max_, cardiac activity and studying effects of oxygen availability during heat ramping. We tested three main predictions: 1) if neural dysfunction in the brain limits thermal tolerance, a neural dysfunction precedes, or coincides with, CT_max_; 2) a seizure or global spreading depolarisation is the neural mechanism that causes CT_max_; 3) neural dysfunction is caused by oxygen limitation and water oxygen manipulation similarly affects both neural function and CT_max_.

## RESULTS

### CT_max_ is similar in larval and adult zebrafish

In Experiment 1, we established a protocol for measuring CT_max_ in five-day-old zebrafish larvae. Activity was recorded in freely swimming larvae in a custom-designed glass heating mantle placed under a camera (**Figure 1A, B**). The larvae were either exposed to a water temperature increasing from 28 °C at a rate of 0.3 °C/min (heat ramp fish, n = 12) or maintained at 28 °C (control fish, n = 14).

**Figure 1.**
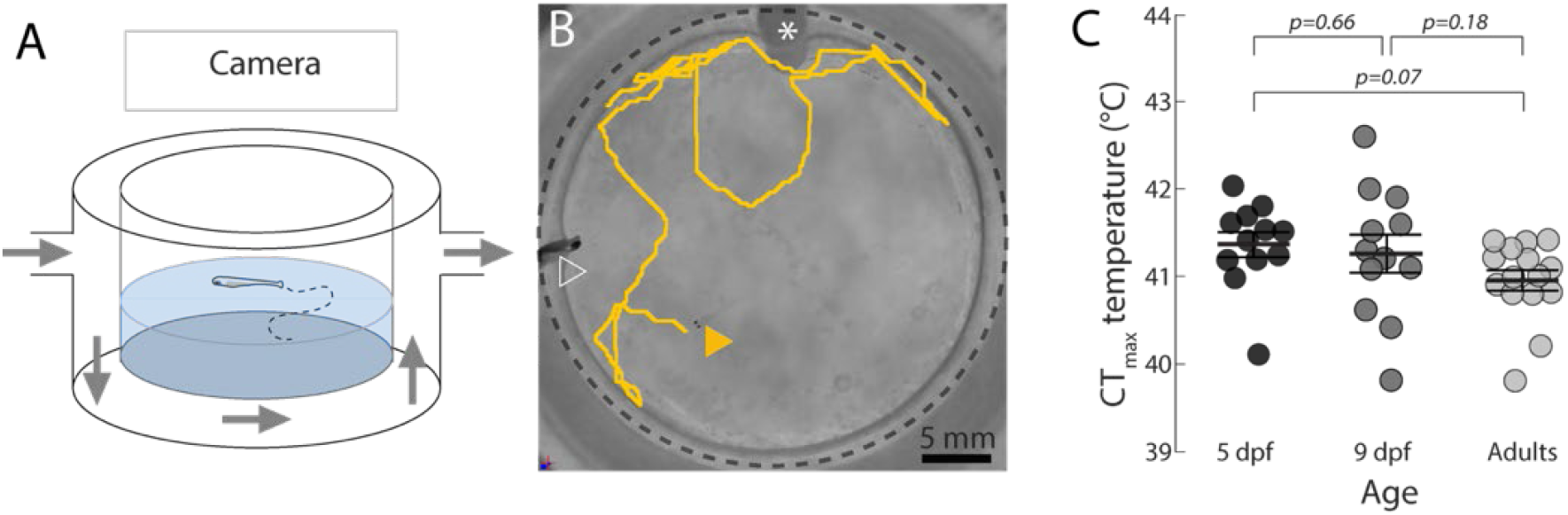
The upper thermal limit in freely swimming zebrafish larvae and adults. Schematic figure (**A**) and photo with top view (**B**) of the experimental setup for CT_max_ measurement in freely swimming zebrafish larvae. **B.** Larva initial position is indicated by a yellow, filled arrowhead and the trajectory is represented in yellow during the following 30 seconds. The dashed black circle indicates the walls of the central chamber. The white arrowhead indicates the position of a thermocouple. The white asterisk indicates the tube for bubbling. **C.** CT_max_ (temperature at loss of response for five and nine dpf and loss of equilibrium for adults, see **Methods**) in five-day-old (n = 12, black circles), nine-day-old (n = 12, dark grey circles) and adult fish (n = 15, light grey circles; LM, nine-day-old: *β* ± S.E. = −0.1 ± 0.2 °C, *t*_(36)_ = −0.5, *p* = 0.7; adult: *β* ± S.E. = −0.4 ± 0.2 °C, *t*_(36)_ = −1.8, *p* = 0.07; *F*_(2,36)_ = 1.9, *R*^2^ = 0.1; **Table S3**). The bars and error bars indicate the group mean and S.E.

To establish a protocol for measuring larval CT_max_, we examined how several activity measures were affected during heat ramping compared to control trials: swimming speed (**Figure S1A**), spiral swimming (**Figure S1B,C; Movie S1**), loss of equilibrium (**Figure S1C, D; Movie S2**), and loss of response (no escape to repeated touches to the trunk, **Figure S1E, Methods**). There was no difference in swimming speed between control and heat ramp fish throughout the assay (**Table S1**). Spiral swimming, loss of equilibrium and loss of response occurred towards the end of the assay in heat ramp fish (**Figure S1B**) and were significantly more prevalent than in control fish (**Figure S1C, D; Table S2, 3**). Yet, spiral swimming was not systematically observed in heat ramp fish, and loss of equilibrium was relatively unspecific, as it also occurred in control fish held at normal temperature. On the other hand, all heat ramp fish eventually became unresponsive to repeated trunk stimulation (loss of response), and all control fish maintained a functional escape response throughout the assay (**Figure S1E**). We concluded that loss of response was a systematic and specific measure of loss of function during warming and therefore a suitable endpoint for determining larval CT_max_.

Using the loss of response as the endpoint for CT_max_, we found that five-day-old heat ramp fish reached CT_max_ at 41.4 ± 0.1 °C (**Figure 1C**). We repeated the experiment with nine-day-old fish and found that they reached CT_max_ at similar temperatures (41.3 ± 0.2 °C, *t*_(36)_ = −0.5, *p* = 0.7, LM; **Table S3**). This indicates that the loss of response criterion is stable during larval development (**Figure 1C**). Furthermore, adult zebrafish tested using the loss of equilibrium protocol (8), reached CT_max_ at similar temperatures (41.0 ± 0.1 °C) to those of larvae (5 dpf - adult: *t*_(36)_ = −1.8, *p* = 0.07; 9 dpf - adult: *t*_(36)_ = - 1.4, *p* = 0.18; **Figure 1C**).

### Warming causes neural dysfunctions near the upper thermal limit

Neurons operate best within a certain thermal range (27, 35, 36). Therefore, we hypothesized that neural dysfunction could explain the heat-induced locomotor impairments and loss of response that we observed at the upper thermal limit of the fish (**Figure 1C, Figure S1E**). To measure neural activity during heat ramping, five-day-old larvae expressing the calcium indicator GCaMP6s in most neurons throughout the brain were mounted in agarose (restrained) in the heating mantle and placed under an epifluorescence microscope in Experiment 2 (**Figure 2A, B**).

**Figure 2.**
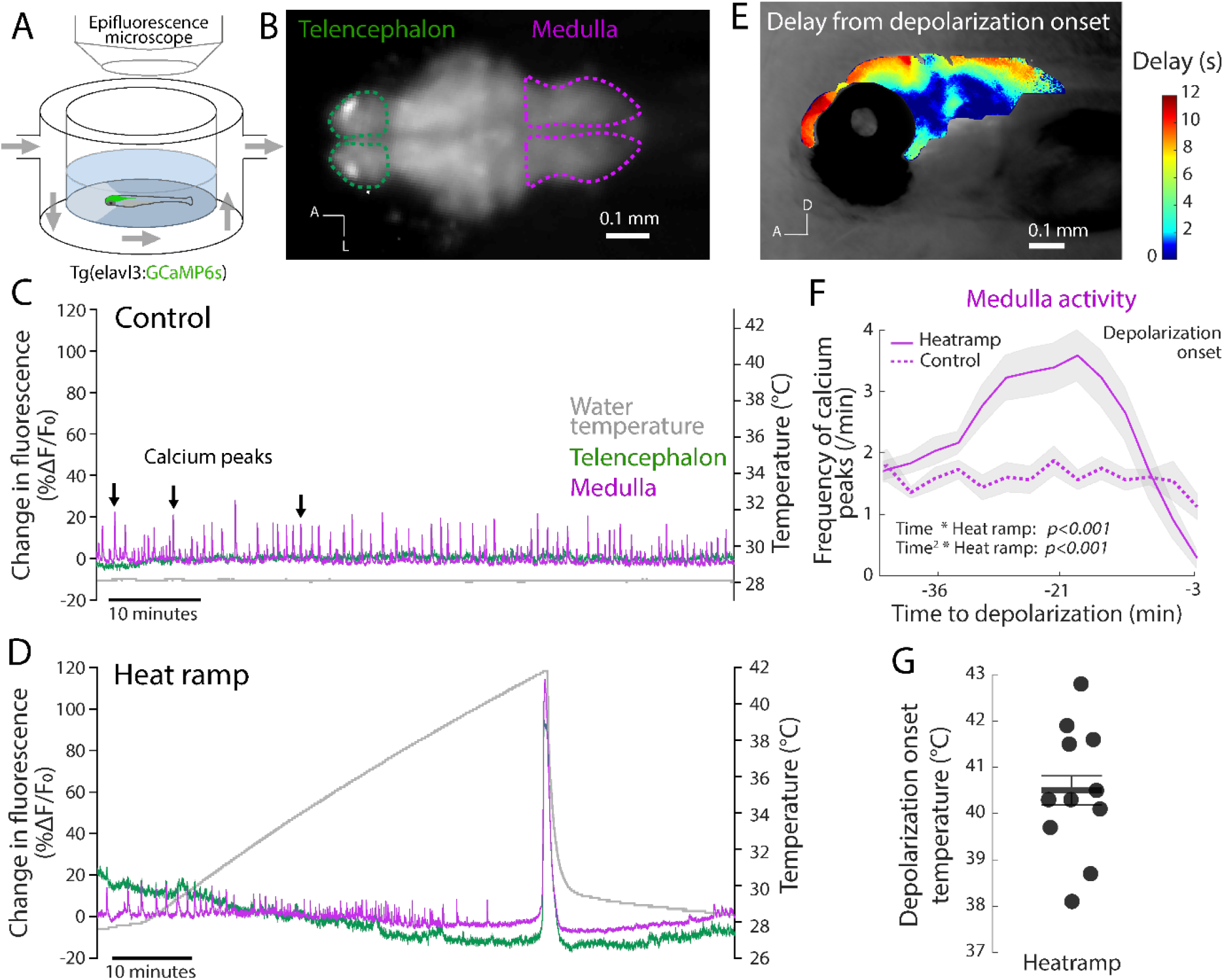
Embedded zebrafish larvae develop brain-wide depolarizations near the upper thermal limit. Schematic overview (**A**) and image (**B**) of the experimental setup for whole brain neural activity measurement in agar-embedded five-day-old *Tg(elavl3:GCaMP6s)* zebrafish larva. **B.** Raw fluorescence image of the larva brain highlighting the telencephalon (dashed green lines) and medulla (dashed magenta lines; A = anterior, L = lateral). **C, D** Change in fluorescence (%Δ*F/F*_0_, left y-axis) in the telencephalon and medulla of a representative control larva (**C**) and a representative heat ramp larva (**D**). The water temperature (grey) was maintained at 28 °C throughout the recording (right y-axis, **C**) for the control fish and was increased during a heat ramp treatment (0.3 °C/min) until a brain-wide depolarization was detected, and then rapidly adjusted to 28 °C until the end of the recording (right y-axis, **D**). Note the return to holding temperature in heat ramp fish and recovery of normal brain activity and calcium peaks 10 - 15 minutes after the brain-wide depolarization in **D. E.** Heatmap illustrating the temporal spread of the depolarization throughout the brain in a representative heat ramp fish mounted laterally (see **Methods**; A = anterior, D = dorsal). **F.** Frequency of medulla calcium peaks in heat ramp fish (n = 11; magenta line) during the 50 minutes preceding the brain-wide depolarization, and during the corresponding period in control fish (n = 8, magenta dashed line, (LMM, heat ramp: *β* ± S.E. = −0.4 ± 0.2, *t*_(41)_ −1.1, *p* = 0.3, time: *β* ± S.E. = 0.0 ± 0.0, *t*_(243)_ = 0.9, *p* = 0.4, time^2^: *β* ± S.E. = −0.0 ± 0.0, *t*_(243)_ = −1.1, *p* = 0.3, time x heat ramp: *β* ± S.E. = 0.2 ± 0.0, *t*_(243)_ = 8.0, *p* < 0.001, time^2^ x heat ramp: *β* ± S.E. = −0.0 ± 0.0, *t*_(243)_ = −8.8, *p* < 0.001, *R^2^* = 0.4, **Table S4**). The depolarization onset in heat ramp fish is indicated by a dashed vertical line. **G.** Temperature at depolarization onset in heat ramp fish (40.5 ± 0.4 °C, n = 11). Data are presented with mean (solid line) and S.E. (shaded area) in **F** and with a bar and error bars in **G**.

Prominent neural activity could be observed in the medulla of both heat ramp (n = 11) and control fish (n = 8; **Figure 2C, D**), which was quantified by calculating the frequency of detected calcium peaks (**Figure 2F**). These calcium peaks likely correspond to attempted tail beats, as reported in previous studies (37, 38). Medullar activity remained stable at 1.6 ± 0.1 peaks/min in the control fish (Time: *t*_(243)_ = 0.9, *p* = 0.4, Time^2^: *t*_(243)_ = −1.1, *p* = 0.3, LMM, **Figure 2C, F, Table S4**). Conversely, in heat ramp fish, the medullar activity more than doubled with temperature, and reached 3.6 ± 0.4 peaks/min, prior to a sharp decline below control levels before a large brain-wide depolarization occurred (Time x Heat ramp: *t*_(243)_ = 8.0, *p* < 0.001, Time^2^ x Heat ramp: *t*_(243)_ = −8.8, *p* < 0.001, **Figure 2D, F**, **Table S4**). Plotting medullar activity across temperature confirmed this pattern (**Figure S2**), with a strong decrease in medullar activity compared to control, occurring at 39–41 °C.

During the massive depolarization in heat ramp fish, the telencephalon and the medulla, which were otherwise weakly co-active (**Figure 2C, D**), both displayed a prominent increase in neural activity (**Figure 2D**). The temporal aspect of the spreading depolarization is illustrated in a laterally mounted fish (**Figure 2E, Movie S3**): it spread relatively slowly, and reached the dorsal brain regions 10–12 seconds after the onset in the ventral diencephalon (**Figure 2E**). The heat-induced brain-wide depolarizations started at 40.5 ± 0.4 °C in agar-embedded fish, which is close to the CT_max_ temperature recorded in freely swimming age-matched fish in Experiment 1 (41.4 ± 0.1°C, **Figure 1E**). Upon detection of the brain-wide depolarization, the water temperature was rapidly decreased to 28 °C and brain activity recovered 10–15 minutes after the brain-wide depolarization (**Figure 2D**).

Together, the results of this second experiment show that several heat-induced neural dysfunctions occur near the upper thermal limit of zebrafish larvae. First, neural activity in locomotor brain centres is strongly reduced. Second, a wave of depolarization propagates throughout the brain at temperatures close to the upper thermal limit of freely swimming conspecifics.

### CT_max_ precedes the brain-wide depolarization

To test whether a brain dysfunction limits thermal tolerance, we sought to establish the relative timing of CT_max_ and the heat-induced neural dysfunctions observed in Experiment 2. To do so, we simultaneously measured CT_max_ and neural activity in freely swimming zebrafish during a heat ramp using epifluorescence imaging in Experiment 3 (**Figure 3A, B**). CT_max_(40.9 ± 0.2 °C) preceded the brain-wide depolarization (41.4 ± 0.2 °C, **Figure 3C**) in all individuals, by 0.5 °C on average (t_(6)_ = 4.5, *p* = 0.004, LMM, **Figure 3D, Table S5**). These results from freely swimming fish show that brain-wide depolarizations were not an artefact due to agarose embedding (**Figure 2**). We concluded that CT_max_ is not preceded by the brain-wide depolarization, but instead occurs during the period of reduced medulla activity.

**Figure 3.**
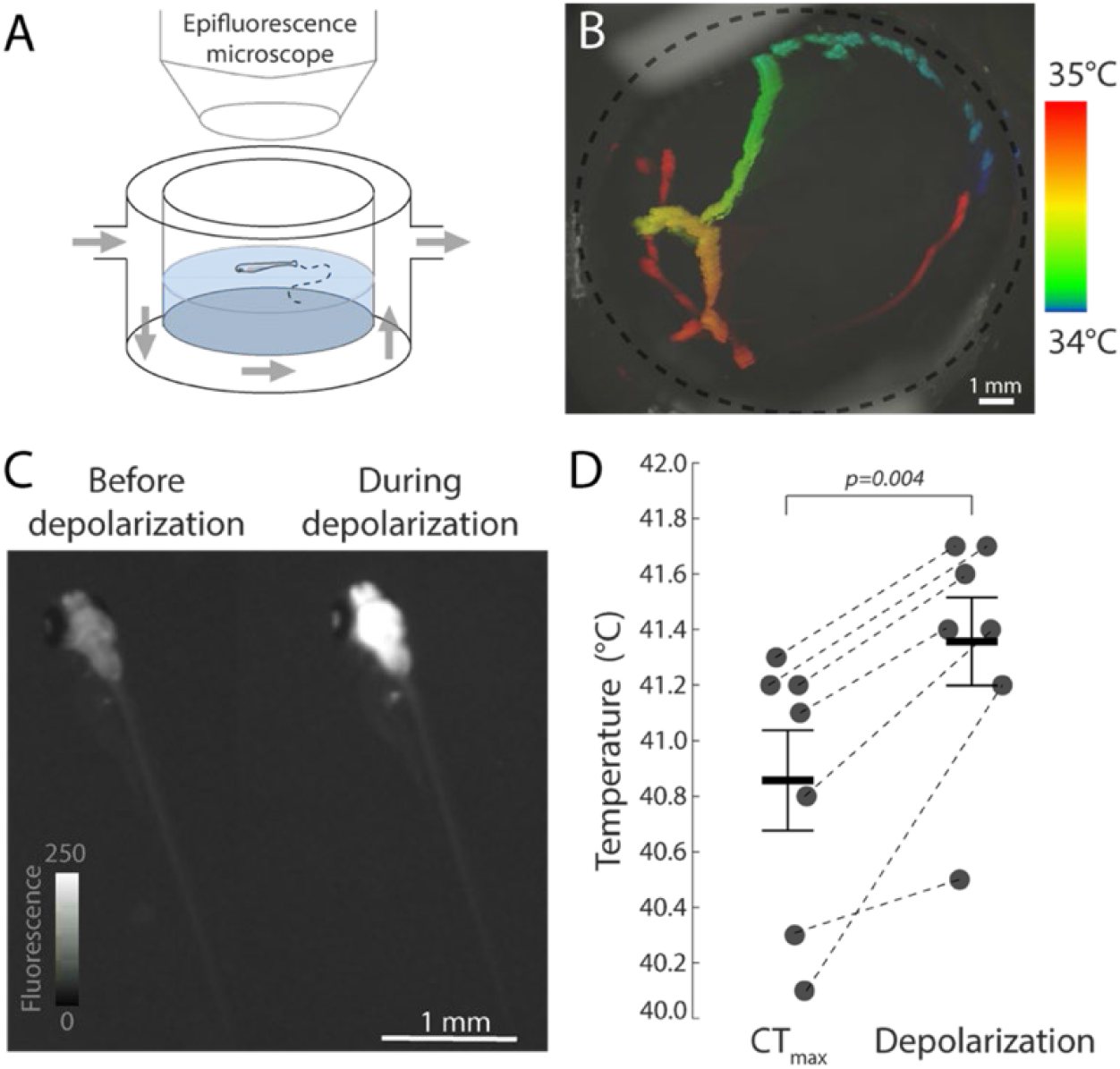
CT_max_ precedes the brain-wide depolarization in freely swimming zebrafish larvae. **A.** Schematic overview of the experimental setup for calcium imaging in freely swimming five-day-old *Tg(elavl3:GCaMP6s)* zebrafish larvae. **B.** Top view of setup with colour-coded time projection of a heat ramp fish’s movements in the setup for three minutes (34–35 °C). **C.** Representative images of brain fluorescence in the same fish as in **B**, before (left) and during (right) the brain-wide depolarization. **D.** CT_max_ and depolarization onset temperatures measured in heat ramp fish (black circles, n=7; LMM, *β* ± S.E. =0.5 ± 0.2 °C, *t*_(6)_ = 4.5, *p* = 0.004, *R^2^* = 0.3, **Table S5**). The bars and error bars indicate the mean and S.E.

### Cardiac function is reduced prior to the brain-wide depolarization

A cardiorespiratory limitation during heating could lead to hypoxic brain tissue and result in neural dysfunctions. To test this hypothesis, we simultaneously measured neural activity and heart rate in larvae embedded laterally in Experiment 4 (**Figure 4A, B**). The heart rate increased with a Q_10_ value (the rate of change with a 10 °C increase in temperature) of 1.5 (from 4.4 ± 0.1 Hz to 5.7 ± 0.1 Hz) between 28 and 34 °C (*t*_(36)_ = 6.9, *p* < 0.001, LMM, **Figure 4C, Table S6**). From 34 to 37 °C, the heart rate decreased (Q_10_ = 0.6) to 4.7 ± 0.2 Hz (*t*_(36)_ = −4.9, *p* < 0.001) after which it remained stable in the period prior to the brain-wide depolarization (*t*_(36)_ = 0.6, *p* = 0.6, **Figure 4C**). The heart rate also remained unchanged during the brain-wide depolarization (**Figure 4D, E**).

**Figure 4.**
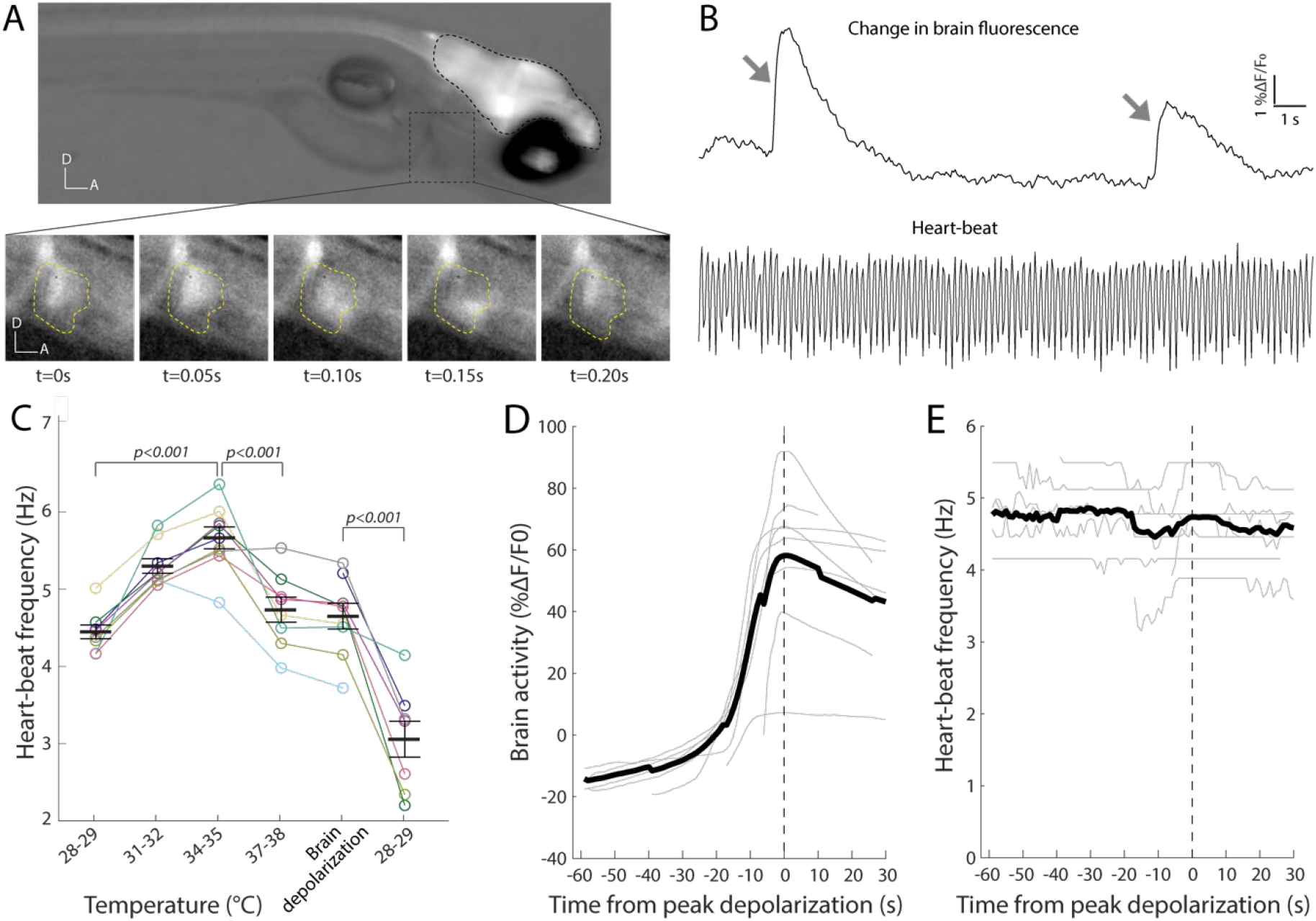
Heart rate before and during the brain-wide depolarization in embedded zebrafish larvae. **A**. Top panel: representative example of simultaneous neural activity and heart rate imaging in a five-day-old *Tg(elavl3:GCaMP6s)* zebrafish larva embedded laterally in agar, using epifluorescence microscopy. The brain is outlined with black dashed lines and the heart with a dashed grey square. Bottom panel: The heart region (dashed yellow line) is shown in inverted greyscale (D = dorsal, A = anterior) during a cardiac cycle. **B**. Top panel: change in brain fluorescence (%Δ*F*/*F*_0_,) in the same fish as in **A**, at 31°C. Change in brain fluorescence are indicated by grey arrows. Bottom panel: change in luminosity within a region of interest in the heart during the same period, which is used to calculate the heart rate. **C**. Average heart rate in embedded heat ramp fish (n = 10). The x-axis shows ranges of 1 °C elevation during which the heart rate was measured. The fifth temperature category (brain depolarization) corresponds to heart rate during the 60 seconds pre- and the 30 seconds post-peak brain depolarization. The sixth category corresponds to heart rate after fish were rapidly returned to holding temperature. Horizontal bars and error bars indicate the mean and S.E. The coloured lines represent individual fish. (LMM: *R*^2^ = 0.8, **Table S6**). Change in neural activity (**D**) and heart rate (**E**) during the brain-wide depolarization (same period as “Brain depolarization” in **C**). Grey lines represent individual fish and the thick black line represent the mean. All traces are aligned with respect to the peak fluorescence (vertical dashed line).

Altogether, no abrupt failure in cardiac activity (e.g., cardiac arrest) was detected during the period prior to the brain-wide depolarization, when CT_max_ occurs. Yet, after an initial increase in heart rate with temperature, the heart rate dropped under further heating. This could lead to tissue oxygen deficiencies, including in the brain, if the cardiac function does not match the increased metabolic requirements at high temperatures.

### Oxygen availability affects neural function and CT_max_

Since the heart rate decreases at high temperatures prior to the brain-wide depolarization (**Figure 4**), it is possible that oxygen deficiency contributes to heat-induced neural dysfunctions preceding CT_max_. In Experiment 5, we tested the effect of oxygen availability on thermal tolerance, by exposing freely swimming larvae to a water oxygen level of either 60 % (hypoxic), 100 % (normoxic) or 150 % (hyperoxic) of air-saturated water. CT_max_ occurred at 39.1 ± 0.1 °C in the hypoxia treatment (n = 24), 0.5 °C lower than in normoxia (39.6 ± 0.1 °C, n = 28, *t*_(73)_ = −2.8, *p* = 0.007, LM, **Figure 5A, Table S7**). Conversely, CT_max_ occurred at 40.1 ± 0.2 °C in the hyperoxia treatment (n = 21), 0.5 °C higher than in normoxia (*t*_(73)_ = 2.3, *p* = 0.025). This indicates that oxygen availability affects thermal tolerance in larval zebrafish.

**Figure 5.**
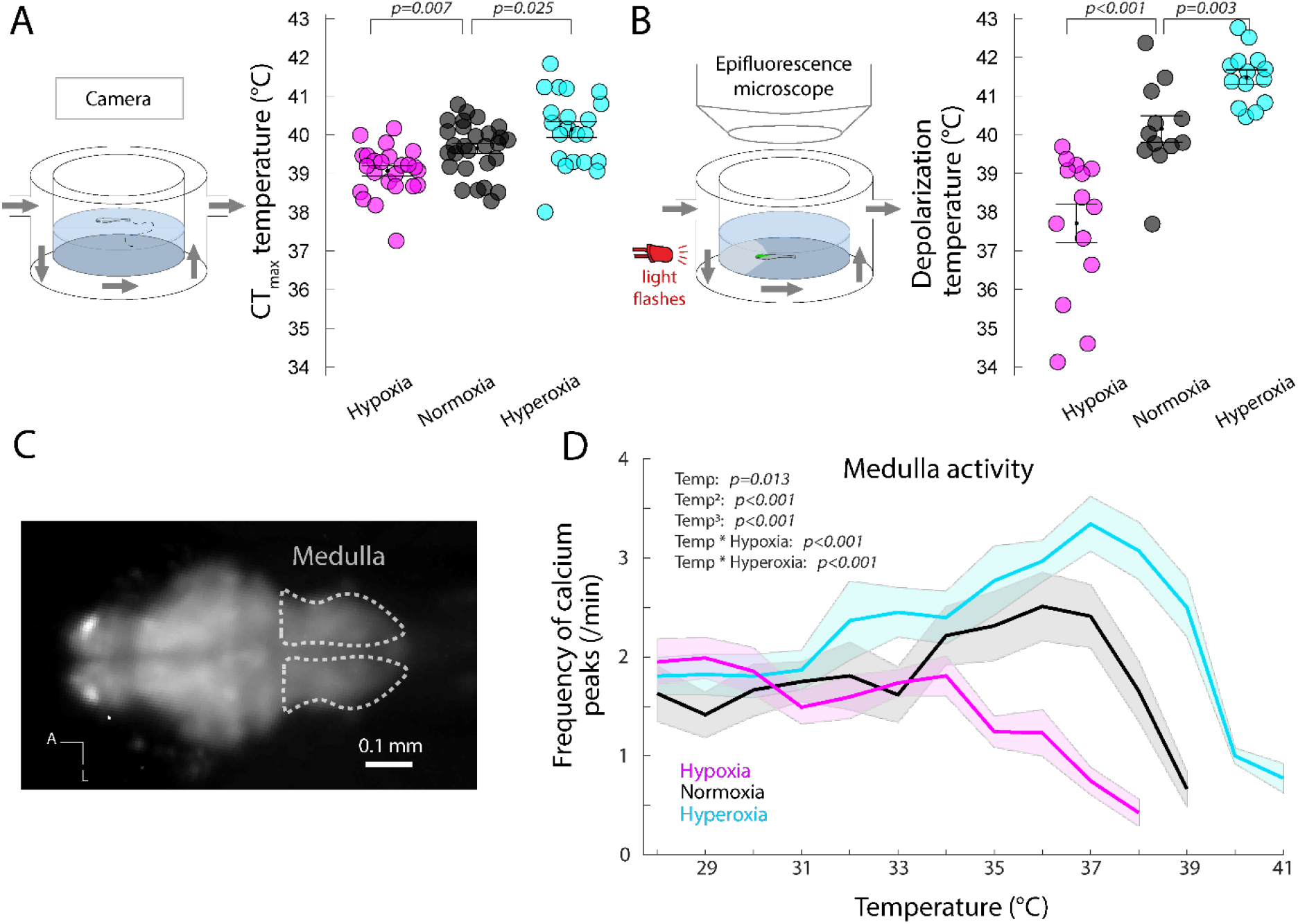
Oxygen availability affects both CT_max_ and onsets of neural dysfunctions. **A.** Effect of oxygen availability on CT_max_ measured in freely swimming fish (setup illustrated in left panel) during heat ramping with water oxygen level of 60 % (hypoxia, magenta circles, n = 24), 100% (normoxia, black circles, n =28), or 150 % (hyperoxia, cyan circles, n = 21) of air saturated water (LM (normoxia) 39.7 ± 0.1; hypoxia *β* ± S.E. = −0.6 ± 0.2 °C, *t*_(73)_ = −2.8, *p* = 0.007; hyperoxia *β* ± S.E. = 0.5 ± 0.2 °C, *t*_(73)_ = 2.3, *p* = 0.025; *F*_(2,70)_ = 11.7, *R*^2^ = 0.3, **Table S7**). **B.** Brain-wide depolarization onset temperatures measured in agar-embedded *Tg(elavl3:GCaMP6s)* five-day-old fish (setup illustrated in left panel) during heat ramping in hypoxia (n = 14), normoxia (n = 12) and hyperoxia (n = 14; LM (normoxia): 41.0 ± 0.3, hypoxia: *β* ± S.E. = −2.7 ± 0.3 °C, *t*_(36)_ = - 6.9, *p* < 0.001; hyperoxia: *β* ± S.E.= 1.2 ± 0.3 °C, *t*_(36)_ = 3.1, *p* = 0.003, Experiment: *β* ± S.E. = −1.8 ± 0.3 °C, *t*_(36)_ = −5.5, *p* < 0.001; *F*_(3,36)_ = 45.5, *R*^2^ = 0.8, **Table S8**). **C**. Image outlining the medulla where the frequency of calcium peaks was measured (A = anterior, L = lateral). **D.** Frequency of medulla calcium peaks during heat ramping in hypoxia (n = 12), normoxia (n = 10) and hyperoxia (n = 11; LMM, normoxia at 28 °C: 1.7 ± 0.2; hypoxia *β* ± S.E. = 0.5 ± 0.3, *t*_(35)_ = 1.8, *p* = 0.09; hyperoxia *β* ± S.E. = 0.1 ± 0.3, *t*_(30)_ = 0.3, *p* = 0.8; temperature *β* ± S.E. = −0.2 ± 0.1, *t*_(355)_ = −2.5, *p* = 0.01; temperature^2^: *β* ± S.E. = 0.1 ± 0.0, *t*_(356)_ = 5.6, *p* < 0.001, temperature^3^: *β* ± S.E. = - 0.01 ± 0.00 *t*_(355)_ = −7.9, *p* < 0.001, hypoxia x temperature: *β* ± S.E. = - 0.2 ± 0.0, *t*_(360)_ = −5.7, *p* < 0.001, hyperoxia x temperature: *β* ± S.E. = 0.1 ± 0.0, *t*_(355)_ = −3.8, *p* < 0.001, *R^2^* = 0.3, **Table S9**). **A, B, D.** Results for hypoxia (60 %, magenta), normoxia (100 %, black) and hyperoxia (150 %, cyan) are presented with mean and S.E. (**A** and **B**: bars and error bars; **D**: solid lines and shaded area).

To test the effect of oxygen availability on neural activity, we also applied the same treatments (60, 100 and 150 % air saturated water) on agar-embedded larvae placed under an epifluorescence microscope. The brain-wide depolarization occurred at 40.2 ± 0.3 °C in normoxia (n = 12) and occurred at 2.5 °C lower in hypoxia (37.7 ± 0.5 °C, n = 14, *t*_(36)_ = −6.9, *p* < 0.001, LM, **Figure 5B, Table S8**). Conversely, the hyperoxia treatment (n = 14) increased the depolarization temperature by 1.3 °C compared to normoxia (41.5 ± 0.2 °C, *t*_(36)_ = 3.1, *p* = 0.003). Oxygen level did not change the amplitude of the brain-wide depolarization, however, the oxygen limitation significantly prolonged the recovery time after the depolarization (**Figure S3A-C, Table S8**). These results indicate that oxygen availability strongly influences the onset of the heat-induced brain-wide depolarization in larval zebrafish.

To further examine the effect of oxygen availability on neural activity, we compared the frequency of calcium peaks in the medulla between the different oxygen treatments. In all groups, the medullar activity sharply decreased in the minutes prior to the brain-wide depolarization (**Figure S3D**), in line with our previous results (**Figure 2F**). Neural activity in the medulla increased until 36 °C (Q_10_ = 1.8) in the normoxia treatment (n = 10) reaching a maximum frequency of 2.5 ± 0.4 peaks/min. The frequency of calcium peaks increased at a higher rate in the hyperoxia treatment (Q_10_ = 1.9, n = 11) and reached a higher maximum frequency of 3.3 ± 0.3 peaks/min at 37 °C. In the hypoxia treatment (n = 12), contrarily, the frequency decreased with temperature from the start of the trial at an accelerating rate (Q_10_ 28–34 °C: 0.8, 34–38 °C: 0.02, **Figure 5D**, LMM, **Table S9**). This demonstrates that increased oxygen availability preserved neural activity in the locomotor centre and that oxygen limitation reduced locomotor-related brain activity near the upper thermal limit of the fish larvae.

Decreased or increased oxygen availability during heat ramping resulted in depressed or improved neural activity in the medulla and decreased or increased thermal limits, respectively (**Figure 5**). These results can be explained by a causal link between heat-impaired neural function and CT_max_, whereby neural dysfunctions at the upper thermal limit culminates into the larvae’s loss of response. An alternative explanation could be that neural function is preserved at high temperatures under low oxygen, but that the decreased neural activity in locomotor centres instead results from less frequent swimming attempts.

To test whether neural responsiveness is affected at high temperatures and co-vary with thermal limits, we measured light-evoked responses in the optic tectum of the same fish used in Experiment 5 during heating (**Figure 5B, Figure 6A, B**). The amplitude of light-evoked responses decreased with increasing temperature in all treatments (**Figure S4A, B**). To compensate for interindividual variation in the light-evoked neural activity, we categorized the individual response to the light flashes as *response* or *no response* (see **Methods, Figure 6C, Figure S4C**). In normoxia (n = 11), the optic tectum responses were initially recorded to 31 % of the light flashes until 31 °C, after which the responsiveness gradually decreased with further heating (**Figure 6C**). In hypoxia (n = 14), the proportion of neural responses decreased from 33 % from the start of the heat ramp. The hyperoxic treatment (n = 13) increased the resilience of sensory response to heating by 5 °C as a responsiveness of 36 % was maintained up to 36 °C before it decreased with further heating (**Table S10**).

**Figure 6.**
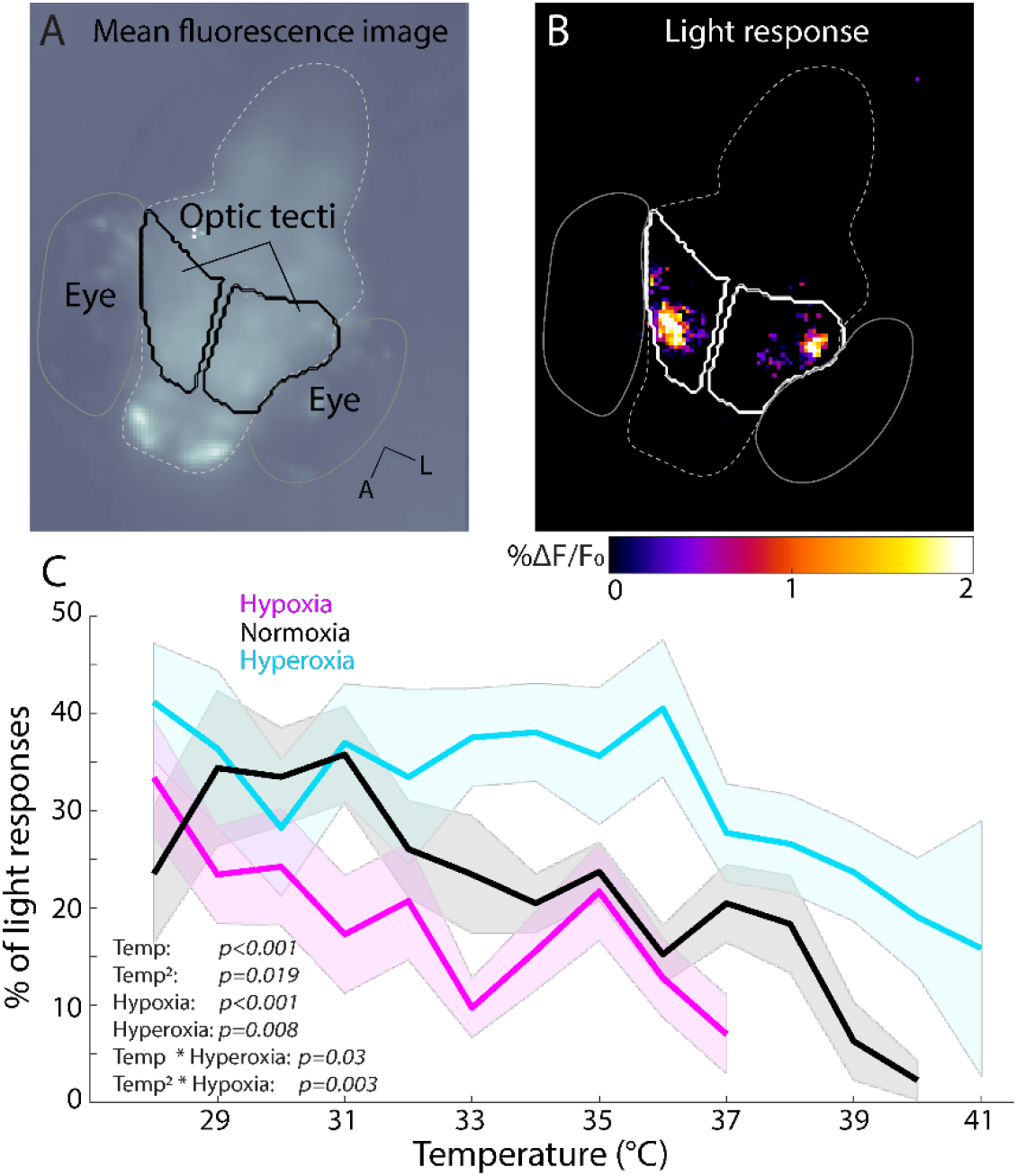
Oxygen availability affects the sensory response resilience to heating. **A**. Mean fluorescence image of the brain (dashed grey line) of an agar-embedded *Tg(elavl3:GCaMP6s)* five-day-old fish under an epifluorescence microscope (same setup as in **Figure 5B**, A = anterior, L = lateral). **B**. Representative light response to a red light flash specifically activating the optic tecti in the same fish as in **A. C**. Percentage of responses to light flashes in the optic tectum during heat ramping with water oxygen level of 60 % (hypoxia, magenta circles, n = 14), 100 % (normoxia, black circles, n = 11), or 150 % (hyperoxia, cyan circles, n = 13) of air saturated water (see **Methods**, GMM, normoxia at average temperature: 1.0 ± 0.1; hypoxia *β* ± S.E. = −0.9 ± 0.2, *t* = −3.9, *p* < 0.001; hyperoxia *β* ± S.E. = 0.5 ± 0.2, *t* = 2.7, *p* = 0.008; temperature *β* ± S.E. = −1.3 ± 0.0, *t*= −5.3, *p* < 0.001; temperature^2^: *β* ± S.E. = - 0.0 ± 0.0, *t* = −2.4, *p* = 0.019, hypoxia x temperature: *β* ± S.E. = 0.1 ± 0.1, *t* = 1.2, *p* = 0.2, hyperoxia x temperature: *β* ± S.E. = 0.1 ± 0.0, *t*= 2.2, hypoxia x temperature^2^: *β* ± S.E. = 0.0 ± 0.0, *t* = 3.0, *p* = 0.003, hyperoxia x temperature^2^: *β* ± S.E. = 0.0 ± 0.0, *t*= 0.5, *p* = 0.6, *R^2^* = 0.8, **Table S10**).

Altogether, these results show that normal neural function is reduced during heating before CT_max_, during the period when freely moving larvae experience locomotor impairments (**Figure S1C**). Moreover, oxygen availability modulated neural activity, the onset of a brain-wide depolarization and CT_max_, suggesting that an oxygen- and heat-induced brain dysfunction sets the thermal limit of larvae zebrafish.

## DISCUSSION

A massive heat-induced global depolarization arose in the larval brain, at similar temperatures as CT_max_ occurred in freely swimming fish. These brain depolarizations were 4–5 times the magnitude of medulla locomotor calcium peaks and lasted considerably longer (**Figure 2**). This abnormally high neural activity in response to heat stress is reminiscent of heat-induced seizures (28–30), and of heat-induced spreading depolarizations (26), which are slow propagating waves of brain depolarization (39). The temporal dynamic of the events measured here is characteristic of the latter. First, the depolarizations spread slowly across the brain (**Figure 2E**), at a speed consistent with that reported for spreading depolarizations (2–9 mm/min, (39, 40)), and slower than seizures (41, 42). Second, the long post-depolarization recovery time with lowered neural activity (**Figure 5E**) is consistent with that seen after spreading depolarizations (39, 43). Overall, the temperatures at which the brain-wide depolarizations occurred, as well as their transient nature, made it a plausible event to explain the reversible unresponsive behavioural state that we observed at CT_max_.

If upper thermal tolerance in zebrafish larvae is caused by a global depolarization shutting down central nervous system function, the depolarization should precede or coincide with CT_max_ measured in the same individual. Surprisingly, freely swimming larval zebrafish reached CT_max_ *before* they developed a brain-wide depolarization. This is in contrast with a series of studies on insects in which measures of upper thermal tolerance match the onset of spreading depolarization in the central nervous system (26, 44). In these studies, upper thermal limits and spreading depolarization onsets were measured in separate individuals, respectively freely moving flies and central nervous system preparations. Using a similar approach in the first part of this article (Experiment 1 and 2), we also found overlapping temperature ranges for CT_max_ and brain-wide depolarization onset. Only when measuring both parameters simultaneously in the same individuals could we resolve the consistent difference between these consecutive events (**Figure 3D**) and determine that CT_max_ precedes the depolarization by half a degree. We thus conclude that brain-wide depolarization is not the cause of CT_max_ in larval zebrafish.

As brain-wide depolarization was found not to be the mechanistic basis of upper thermal limits, we investigated if other forms of neural dysfunction were the cause. We found that neural activity was strongly reduced in locomotor brain regions during the warmest minutes preceding the depolarization (**Figure 2F, 5D, S3D**). Additionally, the neural responsiveness to light stimuli decreased during heating (**Figure 6D**). As both the spontaneous brain activity and the brain response to visual stimulation declined towards the upper thermal limit temperatures, we suggest that gradual brain dysfunction, rather than spreading depolarisation, is the cause for CT_max_.

While these results point to progressively deteriorating brain function causing CT_max_, they are insufficient to determine if the physiological mechanism is through direct thermal effects on neurons, or if the impact is mediated by insufficient brain tissue oxygenation. Therefore, we manipulated oxygen water concentrations to test if oxygen availability alters CT_max_ and neural function. We found that hyperoxia (150 % air saturation) increased both CT_max_ and neural thermal resilience during heat ramping compared to hypoxia (60 %) and normoxia (100 %). Neural activity in the locomotor brain centre and optic tectum response to visual stimuli was more resilient to high temperature in hyperoxia than in hypoxia and normoxia, demonstrating that the thermal impact on the brain is mediated through tissue oxygenation. Furthermore, similarly to the earlier measurements in normoxia, neural activity was largely silenced in the medulla during the very last minutes preceding the brain depolarization, when fish reach their upper thermal limit (**Figure 3D, 5D**). This shows that impairment of the central nervous function coincided with the temperatures where CT_max_ occurs, further suggesting that neural impairment due to lack of oxygen and/or accumulation of anaerobic metabolites contributes to loss of locomotor function observed at the upper thermal limit.

A few studies have found a positive effect of increased oxygen availability on thermal tolerance. Hyperoxia (200 %) improved the upper thermal limit by 1.1 °C in the European perch (45) and by 0.4 °C in the Common triplefin (46). Increased aquatic oxygen availability can extend the survival of several ectotherm species (arthropods, chordates, echinoderms, molluscs, and fish) at extreme temperatures (47, 48). However, hyperoxia did not improve thermal tolerance for a large number of other fish species (23) and only severely hypoxic conditions reduced the upper thermal limit of the red drum and lumpfish (16). Additionally, the aerobic scope, which is the aerobic capacity available for non-maintenance activities, remains high in several ectotherms at thermal limits, indicating surplus oxygen availability at these temperatures (45, 49–52). Taken together these studies suggest that oxygen limitation is not a general mechanism limiting thermal tolerance across all species and contexts. Our results emphasize that interspecific differences in neural tissue resilience to heating and oxygen deprivation, as well as cardiorespiratory capacity might be critical physiological factors explaining these contrasting results between species and contexts.

Cardiac capacity has been suggested to limit thermal tolerance in fish (53), and to test the resilience of the heart we measured heart rate during heat ramping. After the expected initial increase in heart rate as temperature increased (20, 54), the heart rate decreased and then it remained stable upon further heating and through the brain depolarization (**Figure 4C**). Similarly, a decrease or plateau in heart rate with increasing temperatures has been reported for salmonids (18) and European perch (45). The lack of increase in heart rate during heat ramping can contribute to a mismatch between oxygen delivery and tissue oxygen needs. Such a mismatch may cause developing tissue hypoxia and accumulation of metabolites, which can contribute to a loss of response observed at CT_max_. However, five-day-old larvae zebrafish, as in the current experiment, mainly rely on cutaneous respiration (55) and it is therefore unlikely the decline in heart rate had a major contribution to the oxygenation limitation of the brain.

In conclusion, we show that the acute thermal limits of larval zebrafish are not caused by a global brain depolarization but are instead likely caused by a drop in neural activity and responsiveness preceding CT_max_. In accordance with OCLTT predictions, oxygen availability constrains both brain function and the whole animal thermal limits during heat ramping.

## METHODS

### Animals and housing

Experiments were conducted on five-to ten-day-old larvae and seven-to ten-month-old zebrafish (*Danio rerio*). Transgenic zebrafish expressing the calcium indicator GCaMP6s in most neurons (*Tg(elavl3:GCaMP6s)*, (34)) in the *nacre/mitfa* background (56) were used in all experiments. Eggs were collected in the morning and kept at a density of one per mL in fish water (0.2 g of marine salt and 0.04 L AquaSafe per litre of carbon-filtrated water). After hatching at three days post-fertilization (dpf), the larvae were kept in small nursery tanks and after 5 dpf they were fed twice a day with larval food (Tetramin, Tetra). Fish were maintained under standard laboratory conditions (26.8 ± 0.1°C; 12 / 12-hour light / dark cycle). All experimental procedures performed on zebrafish were in accordance with the 2010/63/EU directive and approved by the Norwegian Animal Research Authority (Food and Safety Authority; Permit number: 8578).

### Experimental design

This article consists of five experiments performed on different fish: measuring CT_max_ in freely swimming zebrafish larvae (Experiment 1), recording brain activity during warming in agar-embedded zebrafish larvae (Experiment 2), simultaneous recording of CT_max_ and neural events in freely swimming zebrafish larvae (Experiment 3), simultaneous recording of heart rate and brain activity during warming in agar-embedded larvae (Experiment 4), and recording of CT_max_ and neural function under oxygen manipulation during warming (Experiment 5).

### Measurement of CT_max_

CT_max_ measurements were used in Experiment 1, 3 and 5. All behavioural experiments were conducted in the afternoon, from 12–7 pm.

#### CT_max_ setup

Larval zebrafish swam in the behavioural arena (central compartment) of a double-walled glass heating-mantle made in the NTNU glass workshop. The dimensions were as follows: outer diameter = 42 mm, inner diameter = 29 mm, outer height = 32 mm, inner height = 19 mm. The fish movement was recorded using a webcam (Logitech C270, 720p) positioned above the arena. To enhance contrast, the arena was placed above a background illumination light source. Two outlets connected the glass heating mantle to a water bath. Adjustment of water temperature of the arena was achieved by pumping water from an external heating bath through the heating-mantle. The heating rate in the arena was 0.3 °C per minute following the protocol described in Morgan et al., (8). The water temperature inside the central chamber of the arena was recorded at 1 Hz using two thermocouples (type K, Pico Technology) connected to a data logger (TC-08, Pico Technology).

#### CT_max_ assay in larval zebrafish

The arena was filled with 3 mL of 28 °C water supplied with air bubbling through a modified hypodermic needle fixed to the side wall of the compartment. At the beginning of a CT_max_ recording, a single larva was carefully transferred to the arena. All individuals were given 15 minutes to habituate to the arena before the recording started. The CT_max_ assay lasted up to 50 minutes, during which the temperature within the arena increased from 28 °C by 0.3 °C/min until the larvae reached CT_max_. The temperature was kept constant at 28 °C for recordings of control fish. The fish behaviour was recorded at 4–7 Hz during four intervals: 28–29 °C, 31–32 °C, 34–35 °C and 37 °C–CT_max_. For control larvae, recordings of matching durations were taken at corresponding time points. Methods commonly used to determine CT_max_ in adult fish were unsuitable for 5 dpf larvae zebrafish since the loss of equilibrium criterion (8) also occurred in control fish without heat ramp; and muscular spasms (6) could not be reliably quantified due to the larvae’s small size. Thus, CT_max_ was determined as the temperature at which the animal became unresponsive (11, 26, 57), which was defined as the first of three consecutive touches that did not elicit an escape response (loss of response). In both the control and the heat ramp group, the stimulations were applied to the larva’s trunk throughout the assay, using the tip of a capillary micro-ladder (VWR). In cases where the fish did not escape a stimulation, a second stimulation was applied after a minimum of three seconds. When a fish failed to respond to three consecutive stimulations (CT_max_), it was transferred to 28 °C water and visually monitored. Fish that did not recover normal locomotor activity were rapidly euthanized and were not included in the analysis. This happened in six out of 92 larvae, similar to the mortality rates reported in adult zebrafish in in previous studies (8, 58).

#### CT_max_ assay in adult zebrafish

To compare CT_max_ throughout development, 16 adult *Tg(elavl3:GCaMP6s)* zebrafish, aged seven-to ten-month-old, were tested in the CT_max_ setup described by Morgan et al., (8). The fish were tested in groups of eight fish in a rectangular acrylic tank (25 cm long, 20 cm wide, 18 cm deep) filled with nine litres of water. Individual CT_max_ was measured at the temperature of loss of equilibrium determined as two seconds of inability to maintain postural stability (8). The fish were removed immediately after the criterion was reached. All animals survived the test.

#### CT_max_ assay data analysis

The fish position was manually detected in MATLAB (MathWorks 2018) at 28–29 °C, 31–32 °C, 34– 35 °C, 37–38 °C and during the 1 °C increase preceding CT_max_. The average speed was calculated during these intervals. Loss of equilibrium and spiral swimming events were manually labelled by an experimenter during the 12 minutes preceding CT_max_ in heat ramp fish, and during the corresponding period in control fish. Loss of equilibrium was recorded when the fish tilted to the side for at least one second (**Movie S2**). Spiral swimming event was recorded when the fish rapidly swam in a minimum of two consecutive circles with diameter below 5 mm (**Movie S1**).

### Brain activity during warming in agar-embedded larvae

For the imaging of neural activity in embedded fish in Experiment 2, 4 and 5, a five-day-old larva was embedded at the bottom of the central glass compartment in 2 % low-gelling temperature agarose (Merck). The agarose was carefully removed around the eyes and mouth upon hardening and the arena was filled with 3 mL water. Calcium fluorescence was recorded with a custom-made epifluorescence microscope equipped with a 10x water-immersion objective (UMPLFLN, Olympus), a set of GFP emission-excitation filters (FGL400, MD498, MF525-9, Thorlabs) and a mounted blue LED controlled by a driver (MWWHL4, LEDD1B, Thorlabs). Images were collected at 5 Hz via a custom-written Python script using the Pymba wrapper for interfacing with the camera (Mako G319B, Allied vision).

### Simultaneous recording of CT_max_ and depolarization in freely swimming larvae

In Experiment 3, calcium imaging in freely swimming fish was used to record CT_max_ and brain-wide depolarization simultaneously in the same individuals. The double-walled glass heating-mantle was placed under an epifluorescence microscope (AxioImager, ZEISS) equipped with a megapixel camera (AxioCam 506, ZEISS). The larva was swimming at the centre of the arena, within a 13 mm diameter region delimited with a nylon mesh, to match the limited field of view of the microscope (14 x 14 mm). Time-lapse of fluorescent images were recorded at 5 Hz using the Zen software (ZEN Lite Blue, ZEISS). Since the recordings were made in the dark, the experimenter watched the live recording display of the recording on the computer screen to visually guide the pokes to the fish trunk. The nylon mesh created a small thermal gradient in the arena from the central part to the outer area outside the mesh. During the experimental trials, the temperature was recorded outside the mesh to ensure a full view of the fish. The temperature gradient was quantified in an additional experiment over six heat ramps, by recording the temperature in the arena centre and outside the mesh near the outer wall of the arena with two thermocouples at each location. The gradient was accounted for to calculate the water temperature the fish was exposed to.

### Cardiac function and brain activity during heat ramping in agar-embedded larvae

In Experiment 4, heart rate and neural activity were simultaneously recorded at 20 Hz in laterally mounted larvae, under an epifluorescence microscope (AxioImager, ZEISS) equipped with a megapixel camera (AxioCam 506, ZEISS). Recordings were corrected for drift when needed using Fiji’s Template matching plugin. Whole brain and heart regions of interest were selected in Fiji. Fish whose heart was out of focus during more than two recording intervals (one out of 10 fish), and frames where movement artefacts could not be corrected were not included. Heartbeat frequency during a recording interval was calculated using MATLAB’s continuous wavelet transform function (*cwt*).

### Oxygen effects on CT_max_ and neural function during warming

Due to problems with data storage causing the loss of five recordings of neural activity in the initial data collection in July 2020, we set up an additional round of assays for Experiment 5 in January 2021. The effect of the experimental period was tested for in the statistical analyses (see **Statistical analyses**).

#### CT_max_ assay with oxygen level manipulation

The same CT_max_ setup described for Experiment 1 was used to record the effect of oxygen manipulation on CT_max_ in Experiment 5. The arena was intermittently bubbled with pure oxygen or pure nitrogen to increase or decrease oxygen levels, respectively. The bubbling flow rate was manually adjusted using tubing clamps. A fibre optic oxygen probe (OXROB10, Pyro Science) and a temperature probe (TSUB21, Pyro Science) connected to an oxygen and temperature meter (FireSting O2, Pyro Science) were placed in the chamber to monitor and record the oxygen levels during the assay. The oxygen saturation level was displayed in real time using an oxygen logger software (pyro oxygen logger, Pyro Science) and kept at 150 %, or 60 %, of air saturation during the whole CT_max_ assay by manually adjusting the bubbling intensity.

#### Recording neural activity with oxygen level manipulation

The same epifluorescence calcium imaging setup described for Experiment 2, was used to measure neural activity during heat ramping with oxygen manipulation. For control treatment, the water was bubbled with air through a modified hypodermic needle and maintained at 100% oxygen of air saturation during the trials. To decrease oxygen saturation, the water was bubbled with nitrogen and to increase the oxygen level, oxygen was gently blown on the surface. A bare fibre microsensor (OXB430, Pyro Science) and a temperature probe (TDIP15, Pyro Science) were placed in the chamber to monitor and record oxygen levels. Oxygen levels were manually adjusted to either 150 %, or 60 %, of air saturated water for the high and low oxygen treatments, respectively. Additionally, the larvae were visually stimulated using pulses of light (2 seconds long, every 30 second) using a red LED equipped with a laser line filter (FL05635-10, central wavelength 635 nm, Thorlabs) positioned in front of the fish. The LED was connected to an Arduino board (Arduino mini) and synchronized with the camera using custom-written Python scripts.

### Calcium imaging data analyses

#### In agar-embedded fish

Calcium imaging recordings were corrected for slow drift due to agarose expansion using the Fiji’s (59) Template matching plugin (*Align slices in stack*). The regions of interest corresponding to the whole brain, the telencephalon, the optic tectum and the medulla were manually segmented.

The raw calcium fluorescence signal was calculated by averaging all pixels within a region. To reproducibly detect the brain-wide depolarization onset time across fish, the whole brain raw fluorescence signal was first processed to filter out the calcium events using a 2^nd^ order Butterworth filter (low pass, 0.01 Hz cut-off frequency). The average value and standard deviation of the filtered trace’s time derivative were calculated during the 17 minutes preceding the approximate onset of the depolarization. The depolarization onset was set when the time derivative exceeded the average baseline value by more than five standard deviations.

For calcium peak detection, the fluorescence change was calculated in the left and right medulla using a sliding window of the previous two minutes. Fish in which the signal was weak and noisy due to uneven mounting were not included in the analysis. The calcium peaks were detected automatically using the MATLAB function *findpeaks* and calcium peak frequency (peaks/min) was then averaged across both medullae.

The peak depolarization amplitude (**Figure S3B**) was determined as the maximum returned by the *finpeaks* function after depolarization onset. The recovery time (**Figure S3C**) was the duration between peak depolarization and the return to baseline value (0 %Δ*F/F*_0_). One fish from the hyperoxic treatment was not included as the recordings ended before the fluorescence reached baseline after the depolarization.

For light responses (**Figure 6, Figure S4**), the percentage change in fluorescence in the optic tecti was calculated as %Δ*F/F*_0_ = (*F_t_* - *F*_0_)/*F*_0_ x 100, where *F*_0_ is the averaged fluorescence during the two seconds before a stimulus and *F_t_* the fluorescence at time *t*. The amplitude of the light response was then calculated by averaging the %Δ*F/F*_0_ during the four seconds after the stimulus. To account for the interindividual variation in fluorescence, optic tectum responses were calculated as binary outcomes. The optic tecti were categorized responsive (*response*) if the light response amplitude exceeded two standard deviations above *F_0_*, and otherwise was categorized as not responsive (*no response*).

#### In freely moving fish

Measuring brain-wide fluorescence during the freely swimming assay in Experiment 2 was cumbersome due to the fish moving and tilting. Thus, the depolarization onset temperature was determined by two investigators, who independently selected the time at which the fluorescence increased abruptly in the brain. The final depolarization onset value was obtained by averaging both estimates (agreement between experimenters on average within 19 ± 4 seconds out of 55 minutes).

### Statistical analyses

Data are reported in the text as mean and standard error of the mean (S.E.). All statistical analyses were conducted in R version 4.1.1 (60). We used linear regression models (LM) on independent measurements and the assumptions of normality and homoscedasticity of residual variance were assessed visually using residual plots. Alternative models were considered if assumptions were violated. For responses recorded as repeated measures on individual fish mixed-effects models (LMM) were created using the *lmer* function from the *lme4* package v.1.1-27.1 to account for fish identity as random effect. Data for Experiment 5 were collected over two sampling periods and *Experiment* (1 or 2) was included as a fixed factor in the models to test the effect of this. Akaike information criterion was used to consider interactions in models with two or more predictor variables. Models are also presented with and without data points that were considered extreme outliers (< Q1 - 3 x IQR or > Q3+ 3 x IQR). Summaries of all models are presented in the supplements (**Table S1–S10**) with effect size of predictor variables (ß), their S.E., *t*-values and *p*-values estimated by the respective models and the *R*^2^ for the fixed effects. The significance of effects was considered with a *p*-value criterion of *p* < 0.05.

#### CT_max_ and behavioural response to heat ramping

To analyse the swimming velocity during heat ramping and in the control group a linear mixed-effects model was performed with treatment and time step as predictor variables and the fish identity as random factor (**Table S1**). The proportion of spiral swimming events was analysed using a Chi-square test to handle the lack of variance in the control group (**Table S2**). A Wilcoxon rank sum test was used to test the treatment effect on loss of equilibrium (**Table S2**). To test the difference in CT_max_ throughout development, a linear regression model was used with CT_max_ as a function of the age. The CT_max_ value of one adult fish was considered an extreme outlier and results from models with and without this fish are included in the supplements (**Table S3**).

#### Change in frequency of calcium peaks with temperature

The effect of heat ramping on the frequency of calcium peaks averaged from the medullae in both brain halves was analysed with a linear mixed-effects model with fish identity as a random factor. Temperature was standardized so that the start of the trial was in the intercept of the model (42 minutes before depolarization) (**Table S4**).

#### Temperature of CT_max_ and depolarization

The temperature at which depolarization and CT_max_ occurred in freely swimming fish was compared with a linear mixed-effects model with temperature as response variable, a categorical fixed effect of response type (CT_max_ or depolarization) and fish identity as a random factor (**Table S5**).

#### Change in cardiac function during heat ramping

Effect of temperature on heart rate was tested with a linear mixed-effects model where fish identity was defined as a random factor. The model was releveled to check for differences between the different temperature steps and the *p-*value significance criterion was accordingly reduced to p<0.001 for this analysis (**Table S6**).

#### Effect of oxygen level on CT_max_ and the brain-wide depolarization

The effect of oxygen level on CT_max_ was analysed with a linear regression with CT_max_ as a function of the treatments normoxia, hypoxia and hyperoxia. Two outliers were removed from the hyperoxia group and one from the hypoxia(< Q1-3 x IQR). The model results are presented with and without the outliers (**Table S7**). Experiment was included as a fixed factor in initial models to test if there was an effect of the two data collection periods. The effect of oxygen on depolarization onset, amplitude and recovery time was assessed using the same approach (**Table S8**).

#### Effect of oxygen level on neural activity

The frequency of calcium peaks during heat ramping averaged across the two medullae was analysed with linear mixed-effects model with an interaction between temperature and treatment and fish identity as a random factor (**Table S9**). Temperature was included with quadratic and cubic effects to model the different treatment groups response to temperature, and temperature was standardized so that the starting temperature (28 °C) was in the intercept. The neural response to light stimuli in the optic tecti were tested using a generalized mixed-effects model (GMM) with binomial data (*response, no response*) and the *logit* link function (**Table S10**). During initial observations of the data, complex relationships between the effects of temperature, oxygen and their interaction were detected. Temperature was therefore centered around the mean temperature to model these effects. Fish identity was included as a random factor.

## Acknowledgements

The authors wish to thank Florian Engert and Alex Shier for sharing the *Tg(elavl3:GCaMP6s)* line, Eline Rypdal for fish care, the members of the Animal Physiology section at the Department of Biology for scientific discussions, and Marius Mæhlum for the heartbeat frequency analysis code.

## Authors contributions

AHA, PH, FJ & FK conceived the project. FJ & FK jointly supervised. AHA, PH & FK built the experimental setups. AHA, PH, PK collected the data. FK wrote the software for data acquisition and analyses. AHA, PH & FK analysed the data, AHA, PH and FK prepared the figures and AHA performed the statistical analyses. AHA & FK wrote the manuscript, with contributions from all authors.

## Data accessibility

All datasets from this study and R code for the statistical analyses and figures will be made available in a public repository (Figshare) upon acceptance.

## SUPPLEMENT

### Supplementary figures (Figure S1–S4)

**Supplementary Figure S1.**
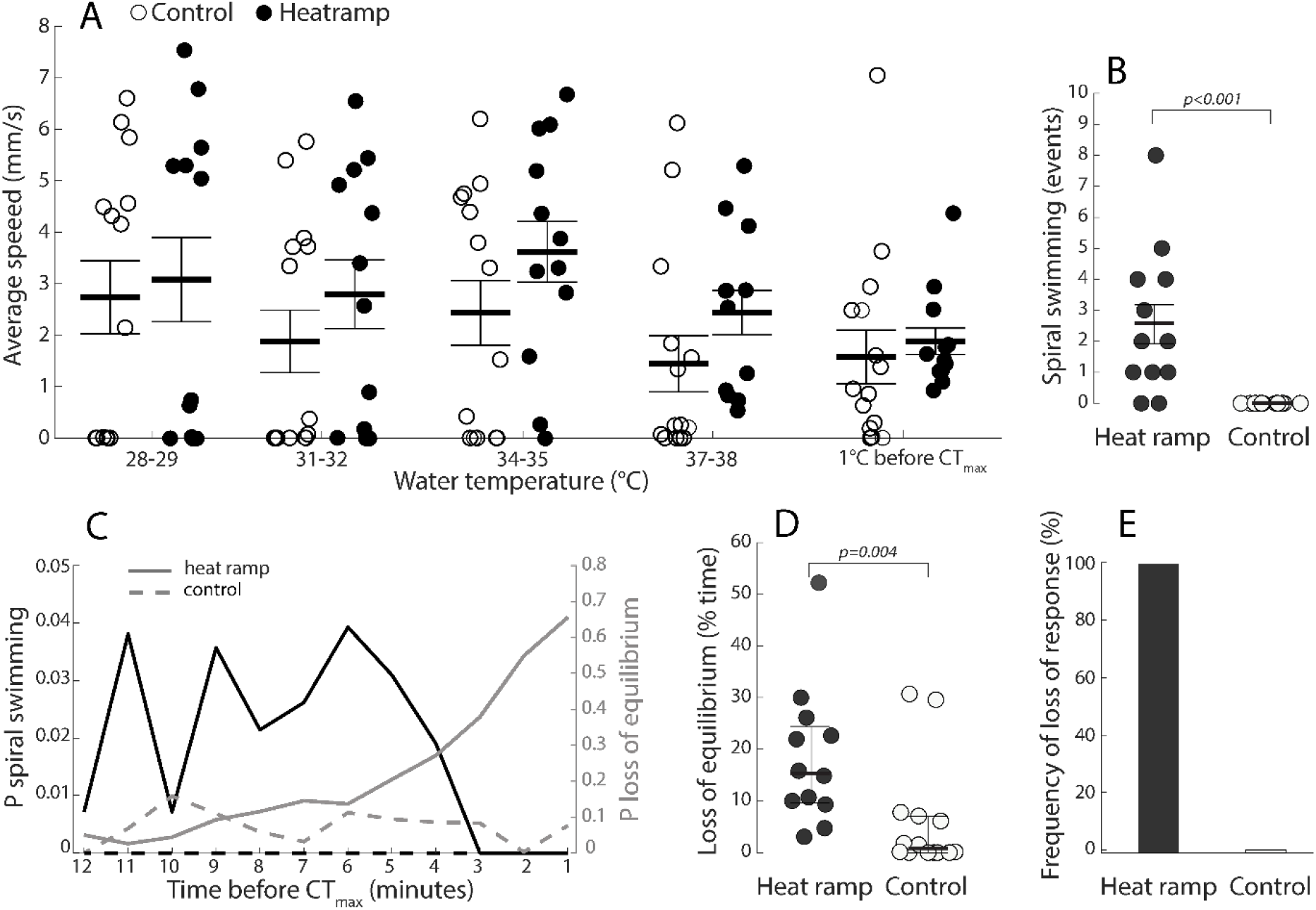
Behavioural responses to heat ramping. **A**. Average swimming speed of five-day-old control (n = 14, empty circles) and heat ramp fish (n = 12, black circles) as a function of temperature. The swimming speed decreased from 2.9 ± 0.4 to 1.7 ± 0.4 mm/s during the assay. There was no difference in the change in swimming speed between the control and the heat ramp treatment (linear mixed-effects model, time (end): *β* ± S.E = −1.2 ± 0.4 mm/s, *t*_(100)_ = −2.4, *p* = 0.015; *R^2^* = 0.1, see **Table S1** for full model output and model including the effect of treatment). **B**. Proportion of spiral swimming events (black), and loss of equilibrium (grey) in control (dashed) and heat ramp fish (solid). **C**. Number of spiral swimming events (see **Methods** and **Movie S1**) recorded during the 12 minutes preceding CT_max_ for five-day-old heat ramp fish (black circles, n = 14), or during the corresponding period in the control fish (empty circles, n = 12; Chi-squared test: *χ*^2^_(df=1,N=26)_ = 15.6, *p* < 0.001, **Table S2**). **D**. Loss of equilibrium (% time, see **Methods** and **Movie S2**) in heat ramp and control fish during the same period as in **C** (Wilcoxon Rank Sum test: *r* = 0.58, *p* = 0.004, **Table S2**). The bars and error bars indicate the group mean and S.E. in **A**. and **C**, and median and inter quartile range in **D. E**. Frequency of individuals that reached the loss of response criterion showing that none of the control fish (n = 14, white bar) and all the heat ramp fish (n = 12, black bar) lost escape response.

**Supplementary Figure S2.**
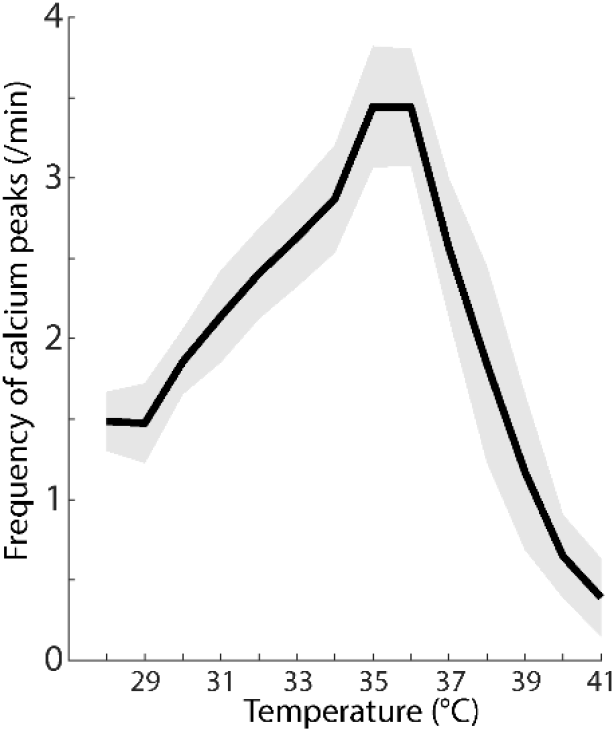
Frequency of calcium peaks during heat ramping over temperature. Frequency of medulla calcium peaks in heat ramp fish (n = 11, same data as displayed in **Figure 2F**) aligned to temperature during the heat ramping. The data is displayed as average (black line) and S.E. (shaded area).

**Supplementary Figure S3.**
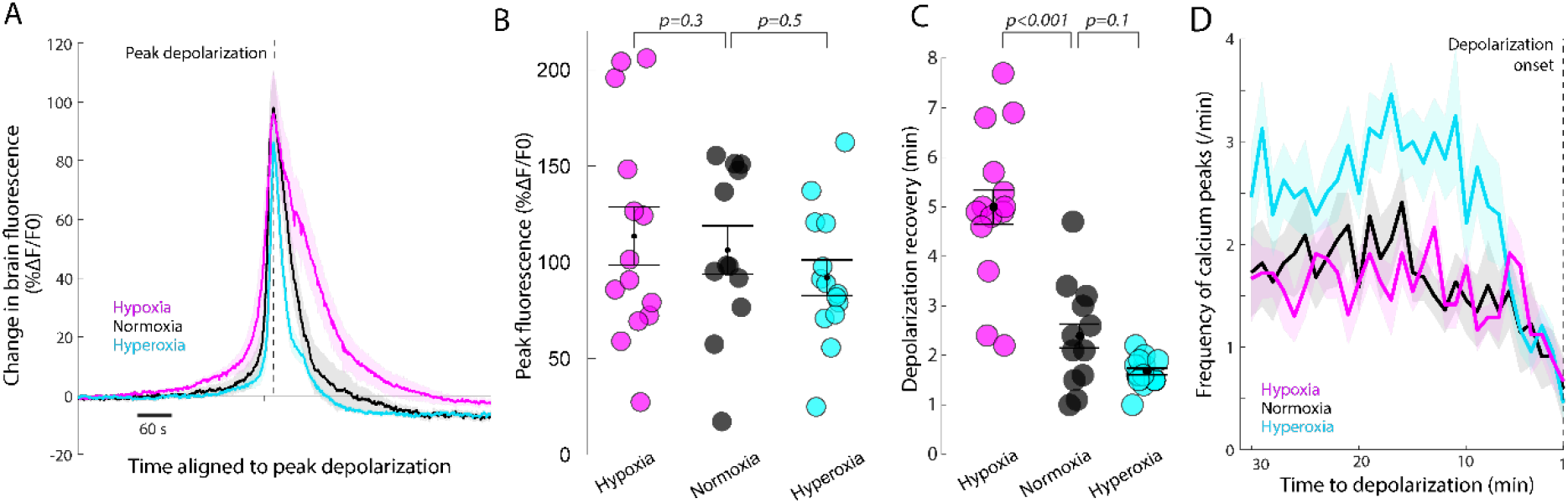
Effect of oxygen level on the temporal dynamics of the brain-wide depolarization. Effects of oxygen availability on medulla activity and the temporal dynamics of the brain wide depolarization. **A**. Change in brain fluorescence aligned to the peak depolarization in hypoxia (n = 14), normoxia (n = 12) and hyperoxia (n = 14). **B**. Amplitude of the brain-wide depolarization in hypoxia (n = 14), normoxia (n = 12) and hyperoxia (n = 14; LM, normoxia: 79.5 ± 12.4 %Δ*F*/*F*_0_; hypoxia: *β* ± S.E. = 14.9 ± 14.8, *t*_(36)_ = −1.0, *p* = 0.3; hyperoxia: *β*±S.E. = −10.6 ± 14.7, *t*_(36)_ = −0.7, *p* = 0.5; Experiment: *β*±S.E. = 53.7 ± 12.0, *t*_(36)_ = 4.5, *p* < 0.001; *F*_(3,36)_ = 7.5, *R*^2^ = 0.3). **C**. Recovery time between peak depolarization and the return to baseline fluorescence in hypoxia (n = 14), normoxia (n = 12) and hyperoxia (n = 13; LM, normoxia: 2.5 ± 0.4 min, hypoxia: *β* ± S.E. = 2.6 ± 0.3, *t*_(35)_ = −1.0, *p* < 0.001; hyperoxia: *β* ±S.E. = −0.7 ± 0.4, *t*_(35)_ = −1.6, *p* = 0.1; Experiment: *β* ±S.E. = −0.4 ± 0.4, *t*_(35)_ = −1.0, *p* = 0.3; *F*_(3,35)_ = 22.8, *R*^2^ = 0.6; **Table S7**). **D**. Frequency of brainstem calcium events recorded during the 30 minutes preceding the brain-wide depolarization in hypoxia (n = 14), normoxia (n = 12) and hyperoxia (n = 14). **A-D.** Results for hypoxia (60 %, magenta), normoxia (100 %, black) and hyperoxia (150 %, cyan) are presented with mean and S.E. (**A, D**: solid lines and shaded area; **B, C**: bars and error bars;).

**Supplementary Figure S4.**
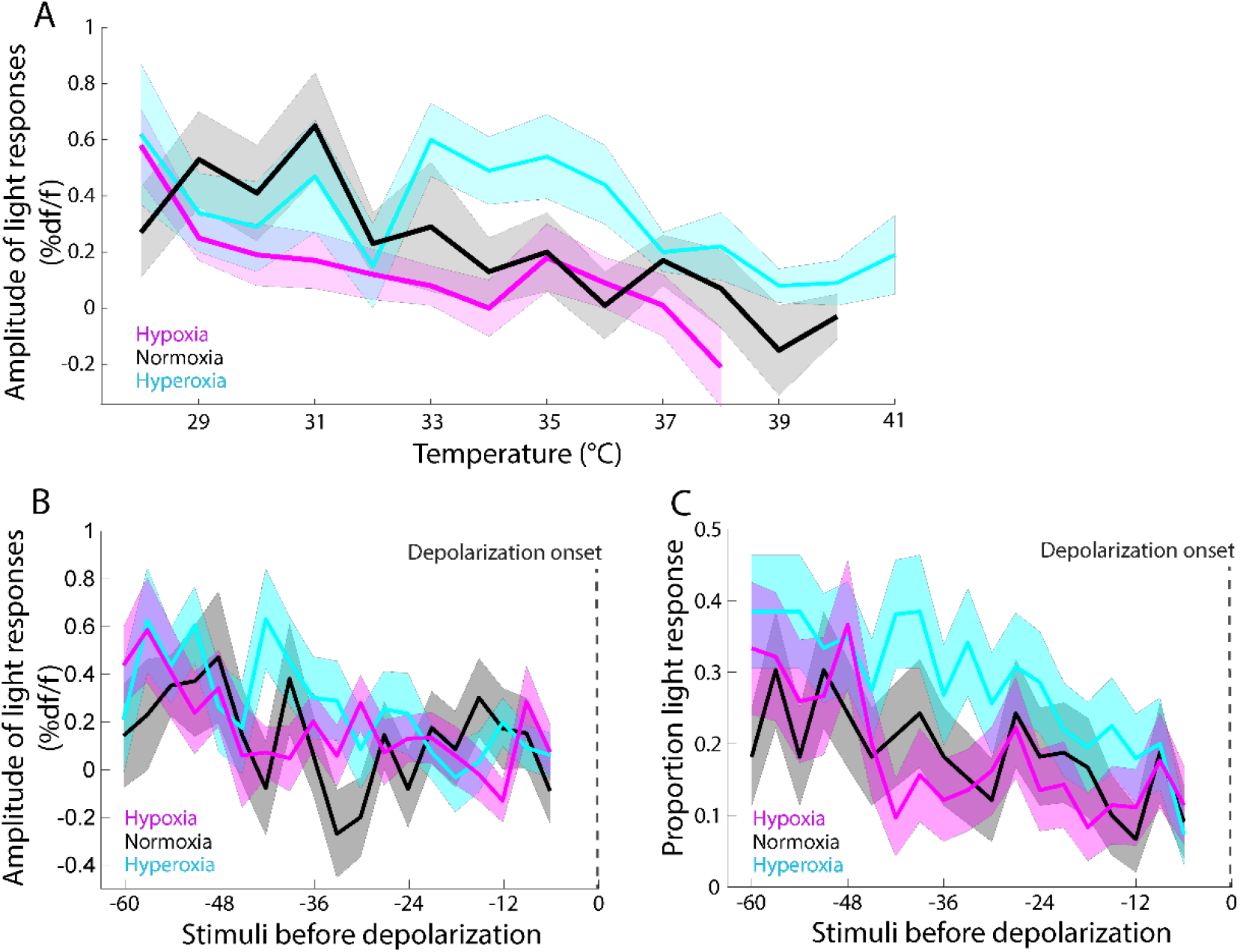
Sensory response in the optic tectum during warming at different oxygen levels. In **A**, the amplitude of the optic tectum response (Δ*F*/*F*_0_) is aligned on temperature during the heat ramping, and in **B**, it is aligned time before depolarization. **C**. Proportion of responses to light flashes in the optic tectum during heat ramping aligned on time at depolarization (same data as in **Figure 6C**). **A-C.** Results for hypoxia (60 %, magenta), normoxia (100 %, black) and hyperoxia (150 %, cyan) are presented with mean (solid lines) and S.E. (shaded area).

### Supplementary tables (Table S1–S10)

**Supplementary Table S1.** Linear mixed-effects model on swim speed. Results of mixed-effects model for swim speed in larval zebrafish (**Figure S1A**) during heat ramping (0.3°C/min) and over an equally long period for control fish (without heat ramping), including fish identity as a random factor. The models include the effect of five time-intervals (activity measured for approximately 3 minutes at five intervals during the trial) and the second model includes the effect of treatment (control or heat ramp). The last measurement for each fish (Time 5) corresponds to the one-degree elevation preceding CT_max_ for the heat ramp fish. The first measurement (Time 1) is in the intercept (and control treatment for the second model) and the units of the estimates are in mm/s.

| *Parameter* | **Swim speed model** |  |  |  | **Model including treatment** |  |  |  |
| --- | --- | --- | --- | --- | --- | --- | --- | --- |
|  | *Estimate ($\beta$)* | *S.E.* | *t-value* | *p-value* | *Estimate ($\beta$)* | *S.E.* | *t-value* | *p-value* |
| Intercept (Time 1) | 2.90 | 0.44 | 6.62 | **<0.001** | 2.55 | 0.52 | 4.88 | **<0.001** |
| Time 2 | -0.59 | 0.48 | -1.23 | 0.222 | -0.59 | 0.48 | -1.23 | 0.222 |
| Time 3 | 0.08 | 0.48 | 0.18 | 0.861 | 0.08 | 0.48 | 0.18 | 0.861 |
| Time 4 | -0.99 | 0.48 | -2.05 | **0.043** | -0.99 | 0.48 | -2.05 | **0.043** |
| Time 5 | -1.18 | 0.48 | -2.43 | **0.017** | -1.18 | 0.48 | -2.43 | **0.017** |
| Treatment |  |  |  |  | 0.76 | 0.62 | 1.22 | 0.233 |
| **Random Effects** |  |  |  |  |  |  |  |  |
| $\sigma^2$ | 3.04 | | | | 3.04 | | | |
| $\tau_{00}$ | 1.94 FishID | | | | 1.89 FishID | | | |
| N | 26 FishID |  |  |  | 26 FishID |  |  |  |
| Observations | 130 |  |  |  | 130 |  |  |  |
| Marginal $R^2$ / Conditional $R^2$ | 0.050 / 0.420 | | | | 0.076 / 0.430 | | | |

**Supplementary Table S2.** Model results for spiral swimming events and loss of equilibrium. Comparisons between the two treatments (heat ramp and control) was done using a Chi-squared test on the proportion of individuals in each treatment that experienced spiral swimming events during the last 12 minutes preceding CT_max_ (0 out of 14 control fish, and 10 out of 12 heat ramp fish). The percentage of time during the same period that the individuals experienced loss of equilibrium was compared with a Wilcoxon rank sum test.

| *Parameter* | **Spiral swimming** |  |  | **Loss of equilibrium** |  |  |
| --- | --- | --- | --- | --- | --- | --- |
| | $\chi^2$ | *df* | *p-value* | *W* | *Estimate (r)* | *p-value* |
| Treatment | 15.6 | 1 | **<0.001** | 27 | 0.58 | **0.004** |
| N = 26: n<sub>control</sub> = 14, n<sub>heat ramp</sub> = 12 |  |  |  |  |  |  |

**Supplementary Table S3.** Linear regression model testing the effect of age on CT_max_. Results for the statistical test for CT_max_ in five-day-old, nine-day-old, and adult zebrafish (**Figure 1C**) testing the effect of life-stage on the upper thermal limit. Model 1 is a linear regression model including all fish whereas in Model 2 one extreme outlier (< Q1-3 x IQR) was removed from the adult group. The five-day-old group is in the intercept and units are in °C.

| *Parameter* | **Model 1** |  |  |  | **Model 2** |  |  |  |
| --- | --- | --- | --- | --- | --- | --- | --- | --- |
|  | *Estimate ($\beta$)* | *S.E.* | *t-value* | *p-value* | *Estimate ($\beta$)* | *S.E.* | *t-value* | *p-value* |
| Intercept (5dpf) | 41.36 | 0.19 | 219.54 | **<0.001** | 41.36 | 0.17 | 250.19 | **<0.001** |
| Age (9dpf) | -0.10 | 0.27 | -0.39 | 0.698 | -0.10 | 0.23 | -0.45 | 0.659 |
| Age (Adult) | -0.54 | 0.25 | -2.16 | **0.038** | -0.41 | 0.22 | -1.84 | 0.073 |
| Observations | 40 |  |  |  | 39 |  |  |  |
| $R^2$ / $R^2$ adjusted | 0.129 / 0.082 | | | | 0.095 / 0.045 | | | |

**Supplementary Table S4.**
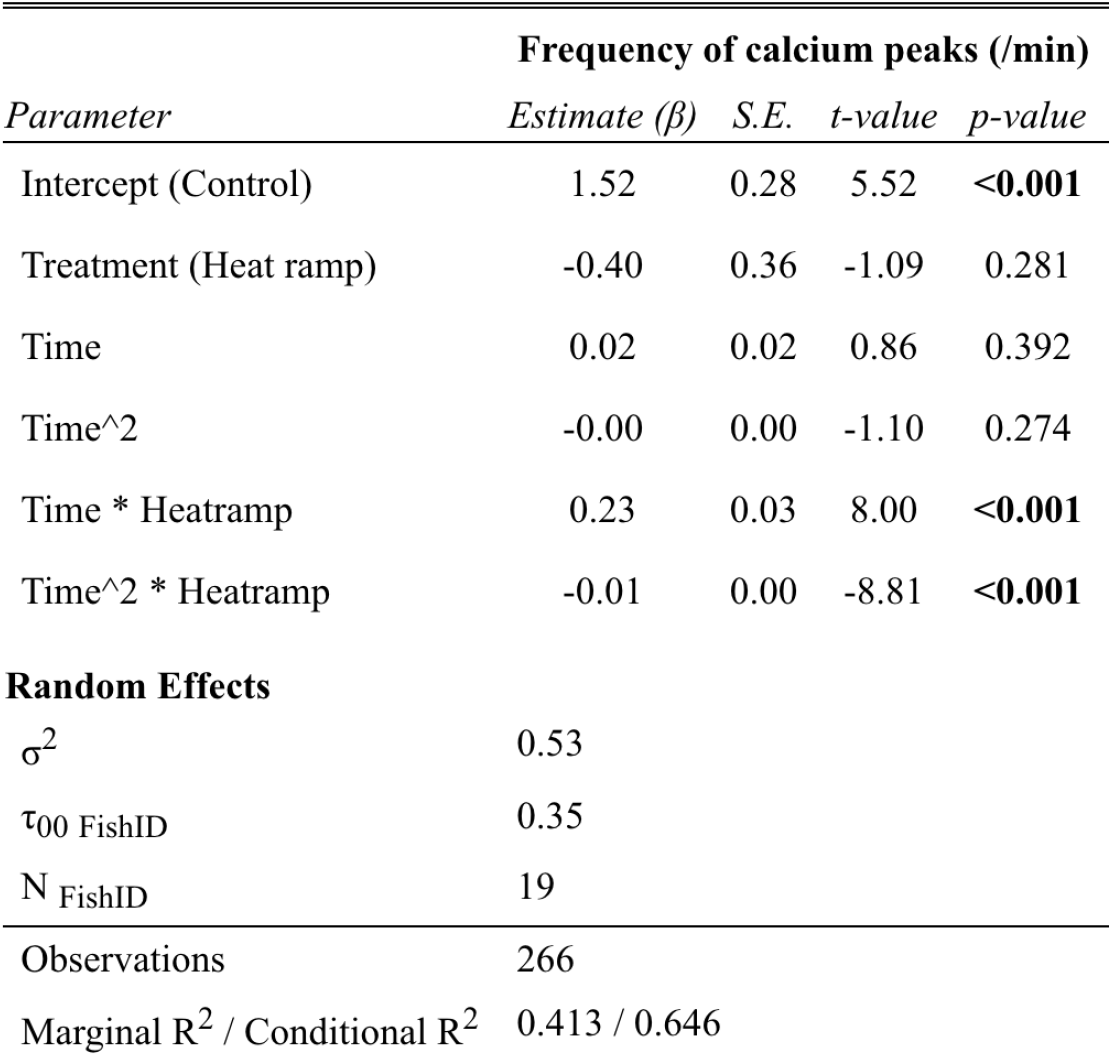
Linear mixed-effects model on the frequency of calcium events during warming. Results on the effect of heat ramping on the average frequency of calcium peaks from the two medullae of larvae zebrafish. The time is standardized so that start of the measuring period is in the intercept. The Time effect is given per minute approaching the depolarization (**Figure 2F**) and includes a quadratic effect to model the thermal effects on the frequency of calcium peaks.

**Supplementary Table S5.** Linear mixed-effects model comparing CT_max_ and depolarization temperature. Results of the statistical test comparing the CT_max_ and brain-wide depolarization temperatures (**Figure 3D**). The mixed-effects model includes fish identity as a random factor. CT_max_ is in the intercept and units are in °C.

| *Parameter* | **CTmax and brain-wide depolarization** |  |  |  |
| --- | --- | --- | --- | --- |
|  | *Estimate ($\beta$)* | *S.E.* | *t-value* | *p-value* |
| Intercept (CTmax) | 40.86 | 0.17 | 240.01 | **<0.001** |
| Brain-wide depolarization | 0.50 | 0.11 | 4.49 | **0.004** |
| **Random Effects** |  |  |  |  |
| $\sigma^2$ | 0.04 | | | |
| $\tau_{00}$ FishID | 0.16 | | | |
| N FishID | 7 |  |  |  |
| Observations | 14 |  |  |  |
| Marginal R<sup>2</sup> / Conditional R<sup>2</sup> | 0.249 / 0.840 |  |  |  |

**Supplementary Table S6.** Linear mixed-effects model on the effect of warming on heart rate. Results of the statistical test for the effect of increasing temperature on heart rate in larval zebrafish (**Figure 4C**) averaged over 1 min intervals during heat ramping. The mixed-effects model includes fish identity as a random factor. The first temperature interval is in the intercept and the units are in Hz.

| Heart rate |  |  |  |  |
| --- | --- | --- | --- | --- |
| *Parameter* | *Estimate ($\beta$)* | *S.E.* | *t-value* | *p-value* |
| Intercept (28°C) | 4.39 | 0.16 | 27.00 | **<0.001** |
| Temperature (31°C) | 0.91 | 0.19 | 4.91 | **<0.001** |
| Temperature (34°C) | 1.27 | 0.19 | 6.88 | **<0.001** |
| Temperature (37°C) | 0.36 | 0.19 | 1.90 | 0.065 |
| Temperature (Depolarization) | 0.26 | 0.19 | 1.40 | 0.170 |
| Temperature (After) | -1.38 | 0.20 | -6.98 | **<0.001** |
| **Random Effects** |  |  |  |  |
| $\sigma^2$ | 0.14 | | | |
| $\tau_{00}$ FishID | 0.07 | | | |
| N FishID | 9 |  |  |  |
| Observations | 50 |  |  |  |
| Marginal $R^2$ / Conditional $R^2$ | 0.753 / 0.836 | | | |

**Supplementary Table S7.** Linear regression testing the effect of oxygen availability on CT_max_. Results of the statistical test for CT_max_ in hypoxia, normoxia and hyperoxia in larval zebrafish testing the effect of oxygen level on the upper thermal limit and the effect of experimental period. Model 1 is a linear regression model including all fish, whereas Model 2 is excluding three extreme outliers (< Q1-3 x IQR), two from the hyperoxia group and one from the hypoxia group. The final model (Model 2.2) is without Experiment as a fixed factor. The normoxia treatment is in the intercept and units for the estimates are in °C.

| *Parameter* | **Model 1** |  |  |  | **Model 2.1** |  |  |  | **Model 2.1** |  |  |  |
| --- | --- | --- | --- | --- | --- | --- | --- | --- | --- | --- | --- | --- |
|  | *Estimate ($\beta$)* | *S.E.* | *t-value* | *p-value* | *Estimate ($\beta$)* | *S.E.* | *t-value* | *p-value* | *Estimate ($\beta$)* | *S.E.* | *t-value* | *p-value* |
| Intercept (Control) | 39.59 | 0.39 | 101.85 | **<0.001** | 39.81 | 0.24 | 164.70 | **<0.001** | 39.65 | 0.14 | 282.63 | **<0.001** |
| Hypoxia | -0.64 | 0.39 | -1.66 | 0.102 | -0.68 | 0.24 | -2.82 | **0.006** | -0.58 | 0.21 | -2.79 | **0.007** |
| Hyperoxia | 0.02 | 0.39 | 0.04 | 0.968 | 0.40 | 0.24 | 1.68 | 0.098 | 0.49 | 0.21 | 2.29 | **0.025** |
| Experiment | 0.06 | 0.34 | 0.17 | 0.862 | -0.18 | 0.21 | -0.85 | 0.399 |  |  |  |  |
| Observations | 76 |  |  |  | 73 |  |  |  | 73 |  |  |  |
| $R^2$ / $R^2$ adjusted | 0.066 / 0.027 | | | | 0.258 / 0.226 | | | | 0.250 / 0.229 | | | |

**Supplementary Table S8.** Effect of oxygen availability on depolarization onset, amplitude, and recovery. Results of the statistical tests on 1) brain-wide depolarization onset, 2) amplitude of depolarization, and 3) recovery time after depolarization, in normoxia, hypoxia and hyperoxia in larval zebrafish (**Figure 5B, Figure S3B, C**) testing the effect of oxygen level on attributes of the brain-wide depolarization. This experiment was conducted in two replicates (different time of year and experimenter and Experiment is included as a fixed effect. Importantly, the treatment effect on depolarization temperature remained strong across the experiments. The normoxic treatment (control) is in the intercept and units are in °C, %Δ*F*/*F*_0_, and minutes, respectively.

| *Parameter* | **Depolarization onset** |  |  |  | **Amplitude of depolarization** |  |  |  | **Recovery time** |  |  |  |
| --- | --- | --- | --- | --- | --- | --- | --- | --- | --- | --- | --- | --- |
|  | *Estimate ($\beta$)* | *S.E.* | *t-value* | *p-value* | *Estimate ($\beta$)* | *S.E.* | *t-value* | *p-value* | *Estimate ($\beta$)* | *S.E.* | *t-value* | *p-value* |
| Intercept (Control) | 41.03 | 0.33 | 125.85 | **<0.001** | 79.48 | 12.35 | 6.43 | **<0.001** | 2.54 | 0.37 | 6.95 | **<0.001** |
| Hypoxia | -2.69 | 0.39 | -6.90 | **<0.001** | 14.88 | 14.80 | 1.00 | 0.322 | 2.57 | 0.44 | 5.86 | **<0.001** |
| Hyperoxia | 1.22 | 0.39 | 3.13 | **0.003** | -10.60 | 14.73 | -0.72 | 0.476 | -0.71 | 0.44 | -1.60 | 0.119 |
| Experiment(2) | -1.76 | 0.32 | -5.55 | **<0.001** | 53.71 | 12.04 | 4.46 | **<0.001** | -0.35 | 0.36 | -0.97 | 0.341 |
| Observations | 40 |  |  |  | 40 |  |  |  | 39 |  |  |  |
| $R^2$ / $R^2$ adjusted | 0.791 / 0.774 | | | | 0.383 / 0.332 | | | | 0.662 / 0.633 | | | |

**Supplementary Table S9.** Model testing the effect of oxygen level on neural activity during warming. Results of the statistical test for frequency of medulla calcium peaks during heat ramping in larval zebrafish (**Figure 5D**) testing the effect of oxygen level (normoxia, hypoxia and hyperoxia) on neural activity. Fish identity is included as a random factor. The model includes temperature with quadratic and cubic effects to model the thermal effects, and an interaction between treatment and temperature. The temperature is standardized so that 28 °C (starting temperature) is in the intercept of the model. The estimates are the average frequency of peaks between the two medullae.

| *Parameter* | **Medulla activity** |  |  |  |
| --- | --- | --- | --- | --- |
|  | *Estimate ($\beta$)* | *S.E.* | *t-value* | *p-value* |
| Intercept (Control) | 1.67 | 0.21 | 8.02 | **<0.001** |
| Treatment (Low) | 0.46 | 0.26 | 1.75 | 0.085 |
| Treatment (High) | 0.07 | 0.26 | 0.28 | 0.782 |
| Temperature (standardized) | -0.19 | 0.08 | -2.50 | **0.013** |
| Temperature (standardized)<sup>2</sup> | 0.08 | 0.01 | 5.64 | **<0.001** |
| Temperature (standardized)<sup>3</sup> | -0.01 | 0.00 | -7.87 | **<0.001** |
| Treatment (Low)*Temperature (standardized) | -0.16 | 0.03 | -5.66 | **<0.001** |
| Treatment (High)*Temperature (standardized) | 0.09 | 0.02 | 3.75 | **<0.001** |
| **Random Effects** |  |  |  |  |
| $\sigma^2$ | 0.49 | | | |
| $\tau_{00}$ FishID | 0.23 | | | |
| N FishID | 33 |  |  |  |
| Observations | 391 |  |  |  |
| Marginal R<sup>2</sup> / Conditional R<sup>2</sup> | 0.318 / 0.533 |  |  |  |

**Supplementary Table S10.** Binomial mixed model on the neural responses to visual stimuli during warming. Effect of oxygen availability on optic tectum response to light stimuli during heat ramping with water oxygen level (normoxia, hypoxia and hyperoxia). Binomial data (response, no response) was tested in a GMM and the results are on *logit* scale. The results can be interpreted in terms of probability of responding to light by using the inverse of the logit function (logit^-1^=*e^x^*/1+*e^x^*). At mean temperature control group had 26 % probability of responding, high had 37 % and low had half of control (13%) (calculated using the inverse logit function).

| *Parameter* | **Light response model** |  |  |  |
| --- | --- | --- | --- | --- |
|  | *Estimate ($\beta$)* | *S.E.* | *t-value* | *p-value* |
| Intercept (Control) | -1.04 | 0.13 | -7.68 | **<0.001** |
| Treatment (Low) | -0.85 | 0.22 | -3.92 | **<0.001** |
| Treatment (High) | 0.49 | 0.18 | 2.67 | **0.008** |
| Temperature (centered) | -0.13 | 0.02 | -5.28 | **<0.001** |
| Temperature (centered)<sup>2</sup> | -0.02 | 0.01 | -2.35 | **0.019** |
| Treatment (Low)*Temperature (centered) | 0.06 | 0.05 | 1.20 | 0.230 |
| Treatment (High)*Temperature (centered) | 0.07 | 0.03 | 2.17 | **0.030** |
| Treatment (Low)*Temperature (centered)<sup>2</sup> | 0.04 | 0.01 | 3.01 | **0.003** |
| Treatment (High)*Temperature (centered)<sup>2</sup> | 0.00 | 0.01 | 0.53 | 0.598 |
| **Random Effects** |  |  |  |  |
| $\sigma^2$ | 3.29 | | | |
| $\tau_{00}$ FishID | 0.06 | | | |
| N FishID | 38 |  |  |  |
| Observations | 2763 |  |  |  |
| Marginal R<sup>2</sup> / Conditional R<sup>2</sup> | 0.080 / 0.083 |  |  |  |

### Supplementary movies (Movie S1-S3)

**Supplementary Movie S1**. Spiral swimming in a five-day-old zebrafish heat ramp larva. Speed 1X.

**Supplementary Movie S2**. Loss of equilibrium in the same heat ramp larva as in Supplementary Movie 1. Speed 1X.

**Supplementary Movie S3**. Representative illustration of the depolarization spreading in the brain of a five-day-old *Tg(elavl3:GCaMP6s)* larva exposed to a heat ramp (same fish as in Figure 2E). Speed 1X.

## REFERENCES

1. S. I. Seneviratne, M. G. Donat, B. Mueller, L. V. Alexander, No pause in the increase of hot temperature extremes. Nature Climate Change 4, 161–163 (2014).

2. J. H. Stillman, Heat Waves, the New Normal: Summertime Temperature Extremes Will Impact Animals, Ecosystems, and Human Communities. Physiology 34, 86–100 (2019).

3. C. A. Deutsch, et al., Impacts of climate warming on terrestrial ectotherms across latitude. PNAS 105, 6668–6672 (2008).

4. P. M. Schulte, The effects of temperature on aerobic metabolism: towards a mechanistic understanding of the responses of ectotherms to a changing environment. Journal of Experimental Biology 218, 1856–1866 (2015).

5. M. L. Pinsky, A. M. Eikeset, D. J. McCauley, J. L. Payne, J. M. Sunday, Greater vulnerability to warming of marine versus terrestrial ectotherms. Nature 569, 108–111 (2019).

6. W. I. Lutterschmidt, V. H. Hutchison, The critical thermal maximum: history and critique. Can. J. Zool. 75, 1561–1574 (1997).

7. C. D. Becker, R. G. Genoway, Evaluation of the critical thermal maximum for determining thermal tolerance of freshwater fish. Environ Biol Fish 4, 245 (1979).

8. R. Morgan, M. H. Finnøen, F. Jutfelt, CT max is repeatable and doesn’t reduce growth in zebrafish. Scientific Reports 8, 7099 (2018).

9. J. M. Sunday, A. E. Bates, N. K. Dulvy, Thermal tolerance and the global redistribution of animals. Nature Climate Change 2, 686–690 (2012).

10. A. Genin, L. Levy, G. Sharon, D. E. Raitsos, A. Diamant, Rapid onsets of warming events trigger mass mortality of coral reef fish. PNAS 117, 25378–25385 (2020).

11. M. J. Friedlander, N. Kotchabhakdi, C. L. Prosser, Effects of cold and heat on behavior and cerebellar function in goldfish. J. Comp. Physiol. 112, 19–45 (1976).

12. F. Jutfelt, et al., Brain cooling marginally increases acute upper thermal tolerance in Atlantic cod. Journal of Experimental Biology 222 (2019).

13. H. O. Pörtner, C. Bock, F. C. Mark, Oxygen- and capacity-limited thermal tolerance: bridging ecology and physiology. J Exp Biol 220, 2685–2696 (2017).

14. H. O. Pörtner, R. Knust, Climate Change Affects Marine Fishes Through the Oxygen Limitation of Thermal Tolerance. Science 315, 95–97 (2007).

15. R. Ern, A mechanistic oxygen- and temperature-limited metabolic niche framework. Philosophical Transactions of the Royal Society B: Biological Sciences 374, 20180540 (2019).

16. R. Ern, T. Norin, A. K. Gamperl, A. J. Esbaugh, Oxygen dependence of upper thermal limits in fishes. Journal of Experimental Biology 219, 3376–3383 (2016).

17. H. O. Pörtner, Climate variations and the physiological basis of temperature dependent biogeography: systemic to molecular hierarchy of thermal tolerance in animals. Comparative Biochemistry and Physiology Part A: Molecular & Integrative Physiology 132, 739–761 (2002).

18. A. P. Farrell, Environment, antecedents and climate change: lessons from the study of temperature physiology and river migration of salmonids. Journal of Experimental Biology 212, 3771–3780 (2009).

19. R. Ern, D. T. T. Huong, N. T. Phuong, T. Wang, M. Bayley, Oxygen delivery does not limit thermal tolerance in a tropical eurythermal crustacean. The Journal of Experimental Biology, 6 (2014).

20. A. P. Farrell, Cardiorespiratory performance during prolonged swimming tests with salmonids: a perspective on temperature effects and potential analytical pitfalls. Philosophical Transactions of the Royal Society B: Biological Sciences 362, 2017–2030 (2007).

21. R. Ern, et al., Some like it hot: Thermal tolerance and oxygen supply capacity in two eurythermal crustaceans. Scientific Reports 5, 10743 (2015).

22. W. C. E. P. Verberk, et al., Does oxygen limit thermal tolerance in arthropods? A critical review of current evidence. Comparative Biochemistry and Physiology Part A: Molecular & Integrative Physiology 192, 64–78 (2016).

23. T. J. McArley, E. Sandblom, N. A. Herbert, Fish and hyperoxia—From cardiorespiratory and biochemical adjustments to aquaculture and ecophysiology implications. Fish and Fisheries 22, 324–355 (2021).

24. F. Jutfelt, et al., Oxygen- and capacity-limited thermal tolerance: blurring ecology and physiology. Journal of Experimental Biology 221 (2018).

25. S. Lefevre, Are global warming and ocean acidification conspiring against marine ectotherms? A meta-analysis of the respiratory effects of elevated temperature, high CO_2_ and their interaction. Conserv Physiol 4 (2016).

26. L. B. Jørgensen, R. M. Robertson, J. Overgaard, Neural dysfunction correlates with heat coma and CT _max_ in *Drosophila* but does not set the boundaries for heat stress survival. J Exp Biol 223, jeb218750 (2020).

27. E. Marder, S. A. Haddad, M. L. Goeritz, P. Rosenbaum, T. Kispersky, How can motor systems retain performance over a wide temperature range? Lessons from the crustacean stomatogastric nervous system. J Comp Physiol A 201, 851–856 (2015).

28. C. Dube, et al., Prolonged febrile seizures in the immature rat model enhance hippocampal excitability long term. Annals of Neurology 47, 336–344 (2000).

29. S. Shinnar, T. A. Glauser, Febrile Seizures. J Child Neurol 17, S44–S52 (2002).

30. R. F. Hunt, G. A. Hortopan, A. Gillespie, S. C. Baraban, A novel zebrafish model of hyperthermia-induced seizures reveals a role for TRPV4 channels and NMDA-type glutamate receptors. Experimental Neurology 237, 199–206 (2012).

31. T. D. Clark, E. Sandblom, F. Jutfelt, Aerobic scope measurements of fishes in an era of climate change: respirometry, relevance and recommendations. Journal of Experimental Biology 216, 2771–2782 (2013).

32. T. D. Clark, E. Sandblom, F. Jutfelt, Response to Farrell and to Pörtner and Giomi. Journal of Experimental Biology 216, 4495–4497 (2013).

33. F. Jutfelt, et al., Response to ‘How and how not to investigate the oxygen and capacity limitation of thermal tolerance (OCLTT) and aerobic scope - remarks on the article by Gräns et al.’ Journal of Experimental Biology 217, 4433–4435 (2014).

34. N. Vladimirov, et al., Light-sheet functional imaging in fictively behaving zebrafish. Nature Methods 11, 883–884 (2014).

35. R. M. Robertson, T. G. Money, Temperature and neuronal circuit function: compensation, tuning and tolerance. Current Opinion in Neurobiology 22, 724–734 (2012).

36. L. S. Tang, A. L. Taylor, A. Rinberg, E. Marder, Robustness of a Rhythmic Circuit to Short- and Long-Term Temperature Changes. Journal of Neuroscience 32, 10075–10085 (2012).

37. L. Turrini, et al., Optical mapping of neuronal activity during seizures in zebrafish. Sci Rep 7, 3025 (2017).

38. M. Ahrens, K.-H. Huang, S. Narayan, B. Mensh, F. Engert, Two-photon calcium imaging during fictive navigation in virtual environments. Frontiers in Neural Circuits 7, 104 (2013).

39. K. E. Spong, R. D. Andrew, R. M. Robertson, Mechanisms of spreading depolarization in vertebrate and insect central nervous systems. J Neurophysiol 116, 1117–1127 (2016).

40. J. Woitzik, et al., Propagation of cortical spreading depolarization in the human cortex after malignant stroke. Neurology 80, 1095–1102 (2013).

41. M. Wenzel, J. P. Hamm, D. S. Peterka, R. Yuste, Reliable and Elastic Propagation of Cortical Seizures In Vivo. Cell Reports 19, 2681–2693 (2017).

42. J. Liu, S. C. Baraban, Network Properties Revealed during Multi-Scale Calcium Imaging of Seizure Activity in Zebrafish. eneuro 6, ENEURO.0041-19.2019 (2019).

43. P. M. Sawant-Pokam, P. Suryavanshi, J. M. Mendez, F. E. Dudek, K. C. Brennan, Mechanisms of Neuronal Silencing After Cortical Spreading Depression. Cerebral Cortex 27 (2016).

44. M. K. Andersen, N. J. S. Jensen, R. M. Robertson, J. Overgaard, Central nervous system shutdown underlies acute cold tolerance in tropical and temperate *Drosophila* species. The Journal of Experimental Biology 221, jeb179598 (2018).

45. A. Ekström, et al., Cardiac oxygen limitation during an acute thermal challenge in the European perch: effects of chronic environmental warming and experimental hyperoxia. American Journal of Physiology-Regulatory, Integrative and Comparative Physiology 311, R440–R449 (2016).

46. T. J. McArley, A. J. R. Hickey, N. A. Herbert, Hyperoxia increases maximum oxygen consumption and aerobic scope of intertidal fish facing acutely high temperatures. Journal of Experimental Biology, jeb.189993 (2018).

47. W. C. E. P. Verberk, R. S. E. W. Leuven, G. Velde, F. Gabel, Thermal limits in native and alien freshwater peracarid Crustacea: The role of habitat use and oxygen limitation. Functional Ecology 32, 926–936 (2018).

48. F. Giomi, et al., Oxygen supersaturation protects coastal marine fauna from ocean warming. Science Advances 5, eaax1814 (2019).

49. A. Grans, et al., Aerobic scope fails to explain the detrimental effects on growth resulting from warming and elevated CO_2_ in Atlantic halibut. Journal of Experimental Biology 217, 711–717 (2014).

50. T. Wang, et al., Anaemia only causes a small reduction in the upper critical temperature of sea bass: is oxygen delivery the limiting factor for tolerance of acute warming in fishes? Journal of Experimental Biology 217, 4275–4278 (2014).

51. J. Brijs, et al., Experimental manipulations of tissue oxygen supply do not affect warming tolerance of European perch. Journal of Experimental Biology 218, 2448–2454 (2015).

52. J. Overgaard, et al., Aerobic scope and cardiovascular oxygen transport is not compromised at high temperatures in the toad *Rhinella marina*. Journal of Experimental Biology 215, 3519–3526 (2012).

53. A. P. Farrell, Cardiorespiratory performance in salmonids during exercise at high temperature: insights into cardiovascular design limitations in fishes. Comparative Biochemistry and Physiology Part A: Molecular & Integrative Physiology 132, 797–810 (2002).

54. M. J. Gollock, S. Currie, L. H. Petersen, A. K. Gamperl, Cardiovascular and haematological responses of Atlantic cod *(Gadus morhua)* to acute temperature increase. Journal of Experimental Biology 209, 2961–2970 (2006).

55. P. Rombough, Gills are needed for ionoregulation before they are needed for O2 uptake in developing zebrafish, *Danio rerio*. Journal of Experimental Biology 205, 1787–1794 (2002).

56. J. A. Lister, C. P. Robertson, T. Lepage, S. L. Johnson, D. W. Raible, nacre encodes a zebrafish microphthalmia-related protein that regulates neural-crest-derived pigment cell fate. Development 126, 3757–3767 (1999).

57. E. Sherman, D. Levitis, Heat hardening as a function of developmental stage in larval and juvenile *Bufo americanus* and *Xenopus laevis*. Journal of Thermal Biology 28, 373–380 (2003).

58. E. R. Åsheim, A. H. Andreassen, R. Morgan, F. Jutfelt, Rapid-warming tolerance correlates with tolerance to slow warming but not growth at non-optimal temperatures in zebrafish. Journal of Experimental Biology, jeb.229195 (2020).

59. J. Schindelin, et al., Fiji: an open-source platform for biological-image analysis. Nature Methods 9, 676–682 (2012).

60. R Core Team, R: A Language and Environment for Statistical Computing (R Foundation for Statistical Computing, 2021).

